# Vocal production differentially affects fast- and broad-spiking neurons in auditory cortex

**DOI:** 10.64898/2026.01.15.699648

**Authors:** Francisco García-Rosales, Boris Sotomayor-Gómez, David Poeppel, Julio C. Hechavarria

## Abstract

Vocalization, foundational for information exchange in animals, is supported by vocalization-modulated neurons that occur across species and brain areas. These neurons are typically studied in an undifferentiated manner, despite the heterogeneous nature of the cortical circuitry regarding cell types. Overlooking this diversity risks producing incomplete models of cortical function. Here, we address this challenge by studying large-scale, cell-type specific auditory cortical activity during vocalization in the bat *Carollia perspicillata*, a vocal specialist. We discovered that vocalisation-related modulatory effects in the auditory cortex, a critical attribute of the underlying circuit, differ starkly between putative inhibitory and pyramidal neurons. During calling, novel cell assemblies and temporal spiking patterns emerge wherein inhibitory cells play a pivotal role. Further, vocalization-specific neural decorrelations affect inhibitory cells most strongly. Together, these results demonstrate that vocalization-related neuronal activity is cell-type specific at the single- and multi-cellular levels of cortical organisation and highlight the importance of inhibitory neurons for auditory cortical dynamics during vocal production.

## Introduction

Vocal production is a fundamental behaviour of a wide variety of vertebrates, including reptiles, birds, and mammals. In the latter, several cortical and subcortical brain structures activate and coordinate to plan ^1–4^, execute ^5–10^, and monitor ^11–15^ call utterance. Immediately following vocalisation, additional brain networks adapt behaviour according to the particular consequences of calling (e.g. bat echolocation ^16,17^). The neural mechanisms supporting vocal production in the mammalian brain are rich and complex, and many aspects remain to be explored and explained.

While (pre-) frontal and motor cortices are associated with the planning and execution of vocal behaviour, the auditory cortex (AC) is crucial for self-monitoring and feedback control ^14,18^. AC neurons exhibit firing rate modulations during calling that range from suppression to excitation, already hundreds of milliseconds prior to vocal onset ^19,20^. Vocalisation-related AC suppression is a consequence of modulatory inputs from motor planning areas which preclude AC activation to self-generated sounds: the so-called “corollary discharge” or “efference copy” signals ^21–24^. Such modulations are crucial for adjusting AC computations to refine vocal-feedback processing ^12,25^.

As in other sensory systems, AC contains a heterogeneity of neuronal types, including inhibitory (interneuron) and excitatory (pyramidal) cells, organised laminarly in standard koniocortical fashion ^26–28^. Interactions across cell types are a cornerstone element of AC function ^29^, including acoustic processing, learning, and attention ^27,30–32^. The AC suppression to self-generated sounds depends heavily on local inhibitory-excitatory dynamics: inhibitory cells are driven by external motor inputs, dampening local activity^21^. While the complementary and interdependent activity of neural types could also be pivotal for vocalisation-related processes, auditory cortical cell-type specific dynamics during vocal production are by and large neglected in the literature. Failing to acknowledge this heterogeneity of the cellular ingredients in the cortical circuitry can lead to an incomplete or misleading understanding of AC function during vocalisation.

Motivated by this basic gap in our understanding, this study addresses vocalisation-related cell-type specific activity in the AC of a vocal specialist: the bat *Carollia perspicillata* ^33–35^. Bats echolocate during navigation and additionally rely on a rich repertoire of communication utterances to interact in complex social environments ^36^. As in other mammals, the bat AC appears to be involved in vocal processes ^17^, but it is unclear whether and how single cells are affected by vocal production. Here, vocalisation-related AC activity was studied in the form of single-cell spiking and local-field potentials (LFPs), while animals vocalised at will. Recordings with Neuropixel probes allowed us to examine large-scale neural populations across cortical columns. We observed clear modulation of neural activity related to vocal production, including suppression and excitation. However, and in line with the heterogeneity of the cortical circuitry, there were glaring differences in the firing dynamics of putative inhibitory and excitatory cells. Inhibitory neurons were disproportionately more likely to be modulated by vocal production, and modulated cells of each type exhibited distinct modulation time-courses. Inhibitory and excitatory cells formed novel cellular ensembles active exclusively during vocalisation, but inhibitory neurons were more likely to associate into assemblies and appeared more relevant for the maintenance of the temporal cohesion of the cortical spiking. Furthermore, vocalisation-specific decorrelations in neural activity most strongly affected putative inhibitory cells. Altogether, our data show that vocalisation-related activity is profoundly cell-type specific at both the single- and multi-cellular levels of cortical organisation.

## Results

Peri-vocal neural activity was studied in three awake *Carollia perspicillata* bats (2 males) while animals vocalised at will. From a total of 16312 acoustic events, neural activity in the form of spikes or LFPs was analysed in relation to 2042 vocalisations (567 echolocation and 1475 communication; **Fig 1**). Vocalisations were selected after careful manual curation, ensuring that a period of at least 500 ms prior to vocal onset (i.e. pre-vocal) was free from acoustic contamination. Example vocalisations (echolocation and communication) are shown in **Fig. 1A**. Call durations ranged from hundreds of microseconds, to tens of milliseconds (**Fig. 1B**), with significant differences between call types (**Fig. 1B**, Wilcoxon rank-sum test, p = 2.46×10^-8^). Call spectral properties are given in **Fig. 1C, D**, with echolocation and communication calls also exhibiting significant differences in peak frequency (p = 1.478×10^-251^). Such differences are in line with well known properties of each call type in this species ^34,37,38^.

**Fig 1.**
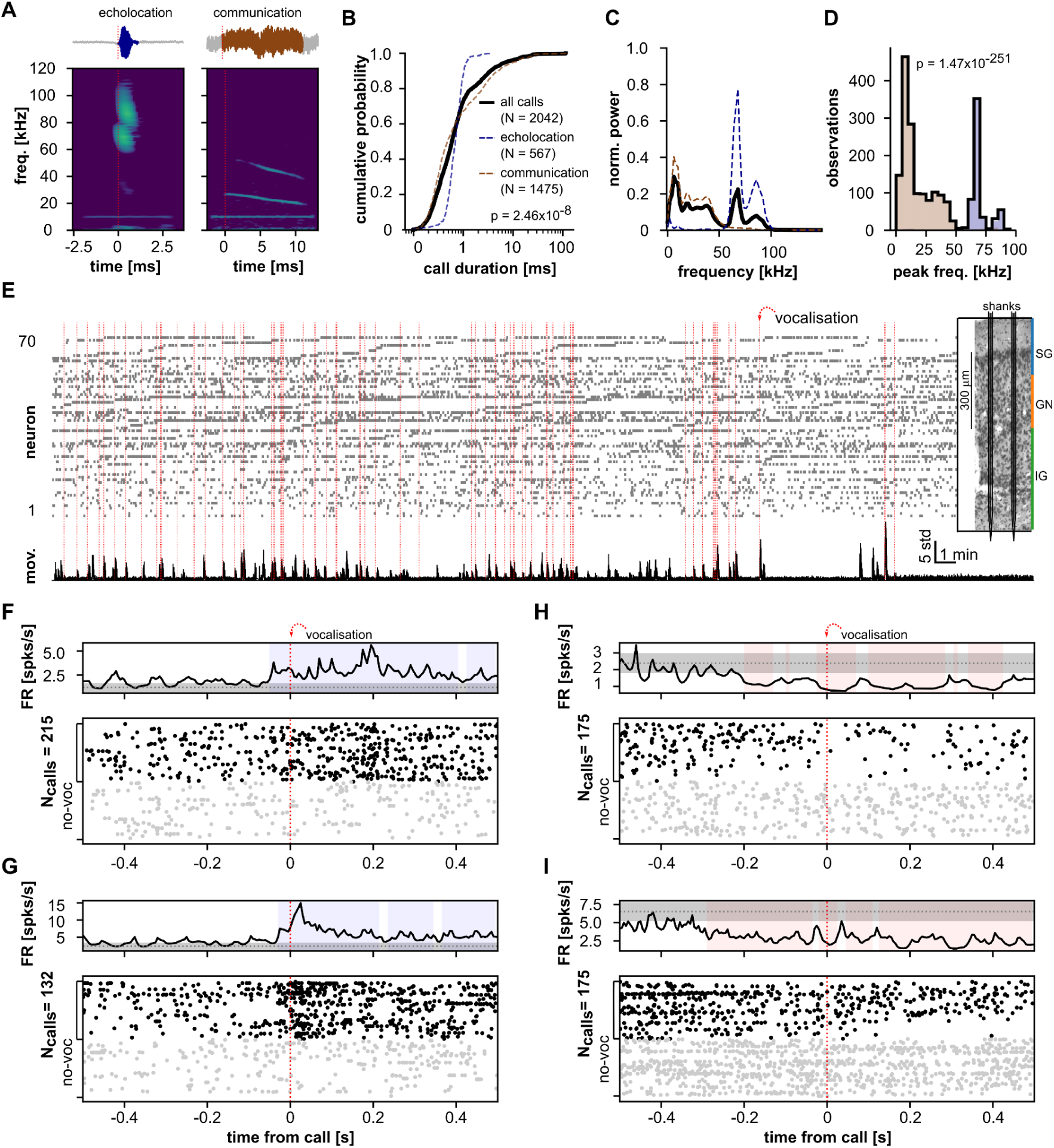
Peri-vocal spiking activity in AC. (**A**) Oscillograms (top) and spectrograms (bottom) of two representative calls from *C. perspicillata*: one echolocation (left) and one communication (right). (**B**) Cumulative distribution of call durations (all vocalisations, black; echolocation, blue; communication, brown). Only vocalisations used for electrophysiological analysis are included. Echolocation calls were significantly shorter than communication ones (Wilcoxon rank-sum test, p = 2.46×10^-8^). (**C**) Average (solid line, mean; shading, s.e.m) spectra of vocalisations. (**D**) Peak frequencies across vocalisations (black outline, all calls; blue shaded area, echolocation; brown shaded area, communication). Values were significantly higher for echolocation calls (Wilcoxon rank-sum test, p = 1.47×10^-251^). (**E**) *Top*: raster plot of simultaneously recorded AC neurons in a representative session; units are sorted according to their depths (spanning 608 µm in total). The inset shows a coronal section of the AC, with two overlaid shanks for illustration, and a depiction of supra-, infra- and granular layer extents (SG, IG, and GN, respectively; neurons are roughly aligned in depth with the anatomy). *Bottom*: animal movement time course. Movement was quantified and z-normalised per session. The movement- and time- scales are given in the figure. Red vertical lines indicate times of vocalisation onsets (only calls used for spiking analysis). For visualisation, a subset of spikes were randomly chosen (N = 5000). (**F**, **G**) Example activity during vocalisation for two excited units. (*Top*) The IFR across vocalisation trials is shown as a solid black line. The 95% confidence interval of no-voc IFRs is depicted as a gray shaded area (mean: gray dashed line). Light blue shaded areas indicate times at which firing rate during vocalisation significantly exceeded spontaneous rates. (*Bottom*) Raster plot corresponding to the IFRs. Spikes belonging to no-voc trials are plotted gray; spikes belonging to vocalisation trials are plotted black. (**H**, **I**) Same as **F**, **G**, but shown for two suppressed units. Light red regions in the firing rates indicate periods where the unit’s firing rate was significantly lower than that observed during no-voc periods.

### Peri-vocal spiking activity in *C. perspicillata*’s AC

**Figure 1E** depicts the activity of 70 simultaneously recorded neurons in a representative recording session. Neural data was concomitantly acquired with animal vocalisations (marked as red lines) and orofacial video, which was used to score the animal’s movement (**Fig. 1E**, bottom trace; see below). Multiple AC units exhibited significant firing rate modulations during vocalisation. These modulations were measured against periods of silence in which animals produced no calls and which were free from acoustic contamination (no-voc periods). Significant modulations were assessed with the use of cluster-based statistical approaches.

The instantaneous firing rates (IFR) of four representative modulated units are shown in **Fig 1F-I**. Two of these units (**F, G**) exhibit significant IFR increase (i.e. excitation) during vocalisation, while the rest (panels **H, I**) exhibit a significant IFR decrease in the vocalisation trial (i.e. suppression). From a total of 592 well-isolated single units with sufficient vocalisation trials (M ≥ 15), 262 (39%) exhibited significant peri-vocal spiking rate modulations. Of these, 123 were excited (47%) and 139 were suppressed (53%; **Fig. S1**). In the majority of these units, significant firing rate modulations began before vocalisation onset. Out of the 123 excited cells, 89 (72%) were already significantly modulated prior to call onset, while 100 (72%) of the 139 suppressed cells also exhibited pre-vocal modulation. The median modulation onset for suppressed units was −200 ms (−377.5–40.5 ms IQR) re. call onset, whereas the median onset for excited cells was −90 ms (−375–0 ms IQR). Most suppression onsets occurred early during pre-voc periods, while the distribution of excitation onsets showed two peaks: one early during pre-voc, and another closer to call onset (see **Fig. S1B**). Median modulation onsets were not significantly different between excitation and suppression (Wilcoxon rank-sum test, p = 0.55), although a 2-sample Kolmogorov-Smirnov test provided marginal evidence that modulation onsets were drawn from different distributions (p = 0.053). Pre-vocal modulation is readily visible in all example units of **Fig. 1**. We also found evidence of modulation throughout the whole peri-vocal period. In such cases, modulation onsets were defined as the earliest possible time in the vocalisation trial (i.e. −500 ms).

### Differential peri-vocal activity for fast-spiking and broad-spiking cells in AC

Auditory cortical units were classified into fast-spiking (FS; putative interneurons) and broad-spiking (BS; putative pyramidal) categories according to their spike waveform shape (see **Fig. S2**). Out of 701 units, 144 (21%) were classified as FS, 511 (73%) as BS, while 46 (7%) were non-classifiable (NC). From all significantly modulated cells (N = 235), 88 (34%) were FS and 147 (56%) BS. These cells were found in all cortical laminae (i.e. supragranular, SG; granular, GN, and infragranular, IG), with most belonging to GN and IG layers (**Fig. 2A**). Neurons in the GN layer were more likely to exhibit suppression than excitation (79 units in total, 49 suppressed, 30 excited; Chi-square test, χ² = 4.57; p = 0.03), which was not the case for IG (162 total units, 84 suppressed, 78 excited; χ² = 0.2; p = 0.03) cells, and only marginally the case for SG (21 neurons, 6 suppressed, 15 excited; χ² = 3.85; p = 0.05) units. The proportion of excitation and suppression, however, did not differ between cell types (FS: 48 suppressed, 40 excited; BS: 80 suppressed, 67 excited; Chi-square test, p > 0.9; **Fig. 2B**). In total, 66% of all analysed FS units (88/134) and 32% of BS cells (147/458) were significantly modulated by vocal production. FS cells were disproportionately more likely to exhibit vocalisation-related modulation (Chi-square test, χ² = 32.75; p = 1.04×10^-8^), suggesting that vocal production affects cell types differentially in the AC.

**Fig 2.**
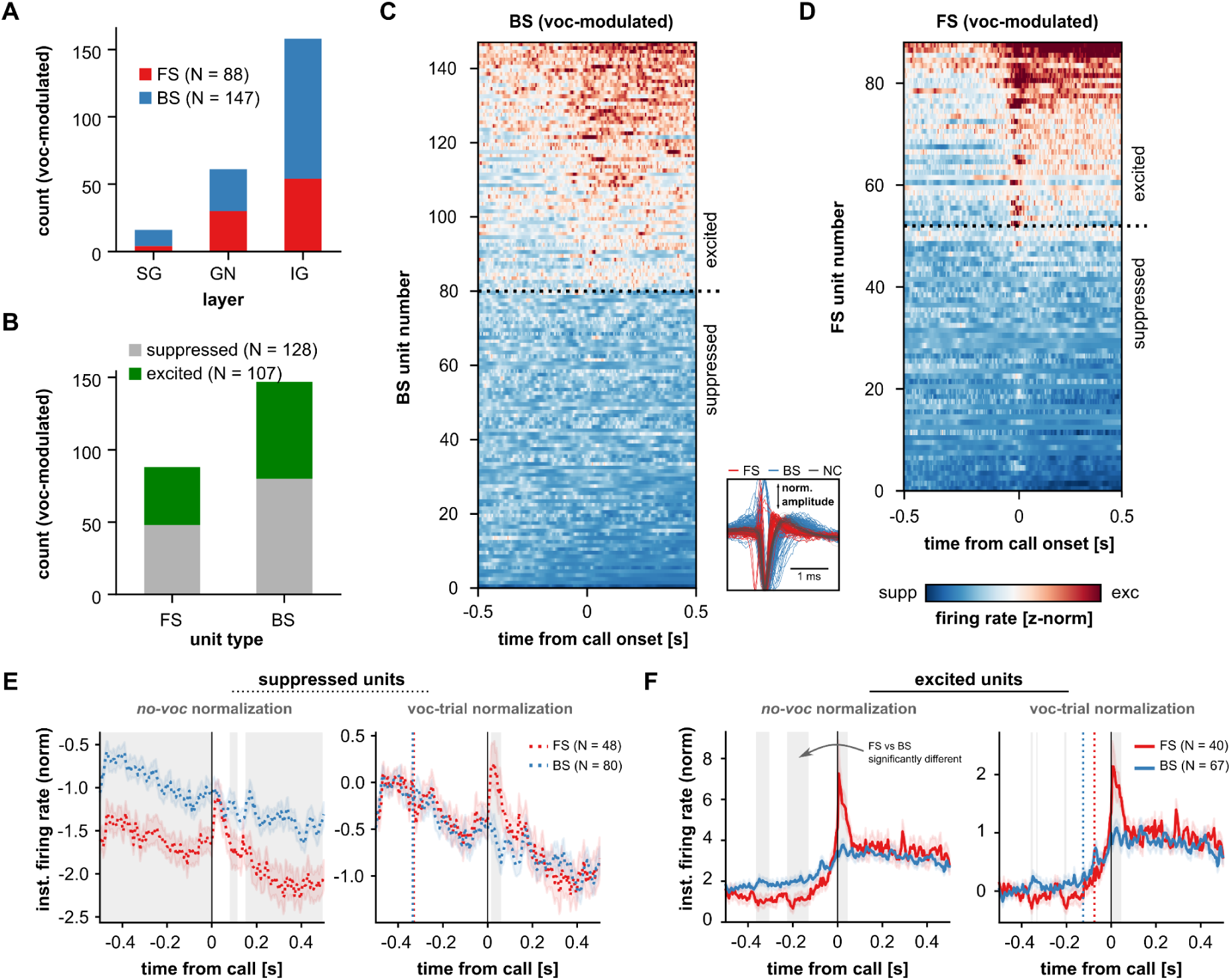
Cell-type specific modulatory dynamics in AC during vocalisation. (**A**) Laminar distribution of significantly modulated units, grouped by cell type (FS: red, BS:blue; SG: supragranular, GN: granular, IG: infragranular). (**B**) Quantification of modulation type (excited: green, or suppressed: gray) for FS and BS cells. (**C**) Peri-vocal IFRs across all significantly modulated BS units (N = 147), sorted according to each unit’s total clustermass (see Methods). (**D**) Same as in **C**, but only for significantly modulated FS units (N = 88). The inset between **C** and **D** depicts representative spike waveforms from FS (red), BS (blue) and non-classifiable (NC; gray) cells. 100 units were selected randomly from the FS and the BS populations for illustration (N_FS-total_ = 144; N_BS-total_ = 511); all 46 NC units were plotted. (**E**) *Left*: Peri-vocal IFRs for suppressed FS (red) and BS (blue) units. Firing rates were z-normalised to no-voc periods. *Right*: the IFRs of each unit were normalised to a baseline period in the vocalisation trial (−475 to −375 ms relative to call onset). (**F**) Same as in E, but data shown for excited FS and BS cells. Gray shaded areas in the figure depict periods where FS vs. BS IFRs were significantly different (cluster-based statistics; see Methods). Vertical dashed lines in B and D (red for FS, blue for BS units) mark the first time point where IFRs deviated significantly from the baseline period.

Next, to further quantify cell-type specific vocalisation-related modulations, we examined differences in the IFRs of FS and BS units. Peri-vocal firing rates were normalised relative to a no-voc baseline, in order to quantify and compare changes relative to spontaneous activity. To capture general vocalisation dynamics, communication and echolocation utterances were pooled together (see **Fig. S3** for call-specific analysis). No-voc-normalised IFRs (**Fig. 2E**, *Left*) revealed significant (FDR-corrected Wilcoxon signed-rank tests, p < 0.05) peri-vocal suppression throughout the whole vocalisation period for both cell types. This suggests the presence of a latent modulation (i.e. a lasting shift in firing rates) during vocal production, potentially related to altered arousal states linked to call utterance (see Discussion). FS and BS units were nevertheless differently modulated: suppression in FS cells was significantly stronger than in BS ones (gray shaded areas in **Fig. 2**). Suppression increased gradually throughout the vocalisation period for both cell types at a seemingly equal rate. To corroborate this observation, we normalised the cells’ IFRs to a baseline period at the beginning (−475 – −375 ms, re. call onset) of the vocalisation trial (i.e. instead of no-voc epochs), thus quantifying not the overall suppression relative to spontaneous activity, but the unfolding of the modulation in the trials. The outcome of this approach is shown in **Fig. 2E** (*Right*). The modulation time-courses of FS and BS neurons overlapped considerably, such that significant differences between the IFRs of FS and BS cell were for the most part abolished. Indeed, the first time point for which the IFR was significantly different to that of the local baseline was almost simultaneous in both cases (FS: −335 ms; BS: −330 ms; vertical dashed lines in **Fig. 2Es**), further confirming the temporal overlap. These results suggest the existence of a common suppressive mechanism for FS and BS, which strengthens over time during vocalisation.

Excited units also exhibited significant, latent modulations throughout the peri-vocal period (**Fig. 2F**, *Left*; p < 0.05). IFRs normalised relative to no-voc epochs were significantly different between FS and BS units already during pre-vocal periods, with the strongest excitation seen in BS cells. However, immediately after call onset, the normalised IFR of FS cells increased phasically, such that FS spiking was significantly stronger than BS spiking between 5–45 ms post call onset. After that period, FS firing dropped and plateaued to the same level of BS firing, while the spiking of both cell types remained relatively elevated for the duration of the vocalisation trial (**Fig. 2F**, *Left*). A normalisation of FS and BS IFRs to a baseline at the beginning of the trial (see above) revealed similar trends, but significant differences across cell types were less obvious (**Fig. 2F**, *Right*). However, in contrast to the case of suppressed units, excited BS neurons deviated on average 50 ms earlier (at −125 ms) from the trial baseline, compared to their FS counterparts (−75 ms).

Sixty-nine of the 235 (29%) vocalisation-modulated neurons were significantly responsive to a battery of natural acoustic stimuli presented to the animals in a passive listening condition (see Methods; **Fig. S4**). FS and BS cells exhibited primary-like auditory responses, with an excitation peak shortly after stimulus onset. We observed no evidence for significant differences in the excitation time-courses between suppressed FS and BS units during passive listening. These results suggest that much of the cell-type specific activity observed during vocalisation is unique to this behavioural condition. Likewise, an analysis of neuronal spiking during high- and low- movement epochs revealed that, although a sizable proportion of AC neurons were modulated by movement *per se*, the neural modulations seen during vocalisation cannot be fully accounted for by movement alone (**Fig. S5, A-I**).

### Novel cell assemblies and temporal firing patterns during vocal production

Beyond single-cell analyses, we examined the mesoscopic organisation of firing patterns across multiple neurons. To that end, we focused first on the formation of cell assemblies during vocal production. Cell assemblies are argued to underlie important computational and cognitive functions, and are considered to be the unit of neural syntax ^39,40^. In the context of this work, the term cell assembly refers to a group of units that exhibit coordinated spiking over time in fixed sequential activation (see ^40^). All units (modulated and not) were included in this analysis.

We detected 206 cellular assemblies across all recording sessions during vocalisation periods. Representative assemblies are shown across three vocalisation trials, for an exemplary recording, in **Fig. 3A** (assembly identity is represented by colour). Assemblies were active throughout the vocalisation trial, but activation peaked at times near the call onset (**Fig. 2B**). Assemblies were predominantly found in GN layers (**Fig. 3C**), with most assemblies having constituent cells across more than one lamina (i.e. trans-laminar assemblies 147/206, 71%). Trans-laminar assemblies were assigned in **Fig. 3C** to the predominant layer of their constituent cells. The maximum distance between constituent neurons was typically in the hundreds of micrometers, with distances being significantly larger in trans-laminar than in mono-laminar assemblies (Wilcoxon rank-sum test, p = 4.34×10^-10^, d = 1.07; Assembly size ranged **Fig. 3D**). Assembly size ranged from 2 to 4 neurons, with the majority of assemblies being constituted by 3 cells (**Fig. 3D**). In terms of assembly composition according to cell type (**Fig. 3E**), most assemblies were composed exclusively of FS units (71/206, 34%), followed by evenly composed assemblies (i.e. equal number of FS and BS cells; 38, 18%), BS only assemblies (15, 7%), predominantly BS assemblies (i.e. mixed assemblies with more BS cells; 5, 2%), and predominantly FS assemblies (1, <0.5%). The remaining assemblies included NC units and were not included in the composition analysis (76, 37%). Remarkably, FS cells constituted exclusively ⅓ of all assemblies, and were present in ∼52.5% of assemblies with classifiable neuron types, although they were a minority in our dataset (21%). A majority of assemblies activated exclusively during vocalisation periods. (They were not present over hundreds of bootstrap repetitions (M = 250) of assembly detection in no-voc periods, nor in the neural activity associated with passive listening.) Over half of the assemblies of any size were vocalisation-exclusive (**Fig. 3E**; proportion indicated in stacked-bars). A similar observation was made when considering assembly composition based on cell types (**Fig. 3F**). These results suggest a reorganisation of AC spiking, leading to the formation of cell assemblies, which is exclusive to vocal production.

**Fig 3.**
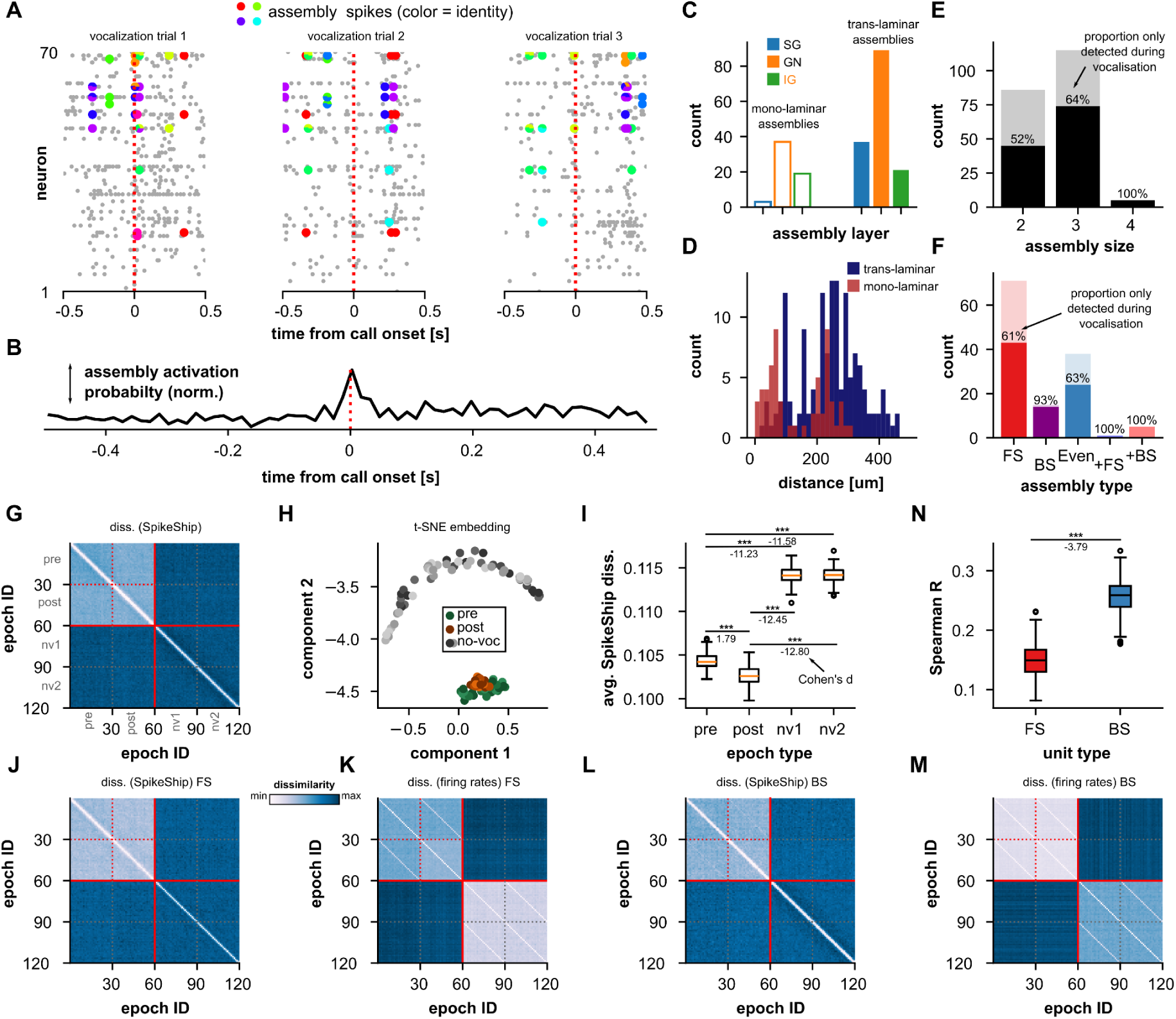
Novel neural assemblies form in AC during vocalisation. (**A**) Representative neural spiking for three vocalisation trials, illustrating the activation of 10 exemplary neural assemblies. Highlighted coloured spikes in the raster plot indicate assembly activation across units as well as assembly identity. (**B**) Time-resolved probability of assembly activation during peri-vocal periods. (**C**) Assembly location according to cortical layers. For trans-laminar assemblies, the most predominant layer is given. (**D**) Distribution of the maximum physical distance between assembly constituent cells. (**E**) Distribution of assembly size (number of constituent cells). Percentages indicate the fraction of assemblies that activate exclusively during vocalisation trials. (**F**) Number of assemblies according to constituent cell types (FS, BS: formed only by FS or BS units; Even: equal number of FS and BS units; +FS, +BS: majority of FS units or BS units, respectively). Percentages as in **C**. (**G**) Median SpikeShip dissimilarity matrix considering all cells. Epoch IDs are sorted according to epoch type: pre-vocal (pre), post-vocal (post) or no-voc (nv) epochs. The diagonal in the matrix was set to the minimum dissimilarity value, for visualization purposes. (**H**) 2-dimensional t-SNE embedding obtained from the dissimilarity matrix in **G**. Colours represent pre-vocal (green shades), post-vocal (orange shades) and no-voc epochs (gray shades). (**I**) Comparison of average dissimilarity for pre-, post-vocal and no-voc epochs across bootstrap repetitions (M = 250), obtained from matrices computed using all recorded units (***, p_corr_ <= 1.97×10^-53^; FDR-corrected Wilcoxon rank-sum tests; effect sizes are indicated as Cohen’s d). (**J, K**) Dissimilarity matrices computed only with FS units. **J**: SpikeShip dissimilarity; **K**: firing rate dissimilarity. (**L, M**) Same as **J,K** but considering BS cells. (**N**) Comparison of correlation between SpikeShip and firing rate dissimilarity matrices for FS and BS units across bootstrap repetitions (***, p = 1.50×10^-81^, Wilcoxon rank-sum test).

To account for potential limitations of the assembly detection approach (e.g., spike-train binning or firing-rate dependance), we applied SpikeShip, a dissimilarity metric based on optimal transport theory ^41^. SpikeShip quantifies the similarity between multi-neuron spiking patterns by computing the minimal transport cost required to align all relative spike-timing relationships across neurons between two epochs. This provides a direct assessment of temporal spiking structure that is not confounded by binning or firing-rate differences. Because this approach gains statistical power from larger populations of neurons ^42^, units were pooled across recordings for the analysis. In total, 417 units with at least 30 vocalisation trials were used (see Methods).

We compared similarity across 30 epochs (one per vocalisation) of pre-vocal and post-vocal activities, as well as two sets of 30 epochs of no-voc periods. To avoid sampling biases, 30 vocalisation trials per neuron were chosen 250 times. The median dissimilarity matrix for 250 iterations is shown in Assembly size ranged **Fig. 3**. A cell *(i, j)* depicts the SpikeShip dissimilarity between epochs *i*, *j*. Lower values indicate more stable temporal spiking patterns across epochs. Relatively low dissimilarity values were observed within (and amongst) pre-vocal and post-vocal epochs (top-left quadrant, **Fig. 3G**), whereas the comparison of any peri-vocal vs no-voc periods yielded higher dissimilarities. The latter suggests that there exists a temporal structure in peri-vocal activity that is clearly distinct from that seen in no-voc periods. A 2-dimensional t-distributed stochastic neighbor embedding (t-SNE) obtained from the dissimilarity matrix in **Fig. 3G** ^41^ illustrates clear separability between spike patterns in peri-vocal periods and their no-voc counterparts (**Fig. 3H**), suggesting that peri-vocal spiking patterns were unique to vocal production. This was also the case when including passive listening epochs in the analysis (**Fig. S6**). Moreover, the vocalisation-related temporal patterns were not fully explained by the animal’s movement alone, as they were not shared with low- or high-movement no-voc epochs (**Fig. S5, J-L**).

The average SpikeShip dissimilarity for pre- and post-vocal epochs across repetitions (M = 250) was significantly smaller than those associated with no-voc periods, with a large effect size (**Fig. 3I**; FDR-Corrected Wilcoxon rank-sum tests, p_corr_ < 3.34×10^-83^; d < −11.23). That is, temporal spiking patterns were significantly more stable during vocalisation. Average dissimilarities were significantly larger for pre-voc than for post-voc epochs with a large effect size (p_corr_ = 1.97×10^-83^; d = 1.79), indicating higher stability of temporal patterns after a vocalisation was produced. The latter could be an effect of acoustic processing in the AC, locked to sensory reafference (but note that this remains vocalisation-specific; see **Fig. S6**). In addition, we examined temporal spike patterns with SpikeShip using exclusively FS (N = 114) or BS (N = 303) units. Similar to when all units were considered, we observed clear separability between peri-vocal and no-voc epochs (**Fig. 3J, L**, left matrices). Altogether, these data support the notion that novel temporal spiking patterns in the AC emerge during vocal production.

The temporal patterns formed by neurons during vocalisation or no-voc periods could be affected by firing rate modulations (see **Fig 2**). To quantify this, we computed dissimilarity matrices using the neurons’ firing rates (see ^42^). Firing rate dissimilarity matrices are shown in **Fig. 3K, M** for FS and BS cells, respectively. Notice that differences in firing rates between peri-vocal and no-voc periods (i.e. vocalisation-related modulations) are echoed in these matrices. Dissimilarity values across epoch types (i.e. pre- or post-vocal vs. no-voc) in the matrices indicate different firing rate dynamics for peri-vocal and no-voc periods across neurons, which are nevertheless relatively stable within epoch types. We quantified the relationship between temporal patterns and firing rate by correlating (Spearman R) the SpikeShip- and the spike-count dissimilarity matrices across repetitions ^42^. Low correlations denote independence between temporal spiking patterns and firing rate. For FS cells, correlation values were relatively low (median R = 0.15; 0.13–0.17 IQR), and were higher for BS neurons (median R = 0.26; 0.24–0.27 IQR). FS-related correlations were significantly smaller than BS-related ones, with a large effect size (Wilcoxon rank-sum test, p = 1.50×10^-81^; d = −3.79; **Fig. 3N**). Thus, FS cells provide temporal information largely complementary to firing-rate modulations, and significantly more so than their BS counterparts.

### Neural desynchronization during vocal production in the AC

In addition to multi-cellular activity patterns, LFPs provide a mesoscopic read-out of neural activity. LFPs constitute a weighted sum of the currents involved in the electrophysiological processes of a large population of neurons, including spiking, synaptic potentials, and other sub-threshold dynamics ^43^. They are considered to be relevant for a variety of neural phenomena including the modulation of local spiking activity, attentional processes, or distant neural communication ^44–46^. Below, we examine LFPs across AC layers and their relationship to vocalisation with a focus on pre-vocal periods and cell types.

Representative peri-vocal LFPs from two different recordings, averaged across vocalisation trials, are given in **Fig. 4A**. Mean pre-vocal and no-voc LFP spectra across recordings are shown in **Fig. 4B**. Low-frequency pre-vocal LFP power was lower than that observed for no-voc periods, independently of the layer being considered. A z-normalisation of the vocalisation-related spectrum using no-voc periods revealed strong suppression of LFP power prior to call onset (**Fig. 4C**). We deemed consistent z-normalised power deviations from 0 across recordings as evidence of significant LFP power suppression in the data. Indeed, for LFP frequencies up to 28 Hz (horizontal bars in **Fig. 4B**), z-normalised power values were significantly different than 0 in all layers (FDR-corrected Wilcoxon signed-rank test, p_corr_ < 0.05).

**Fig 4.**
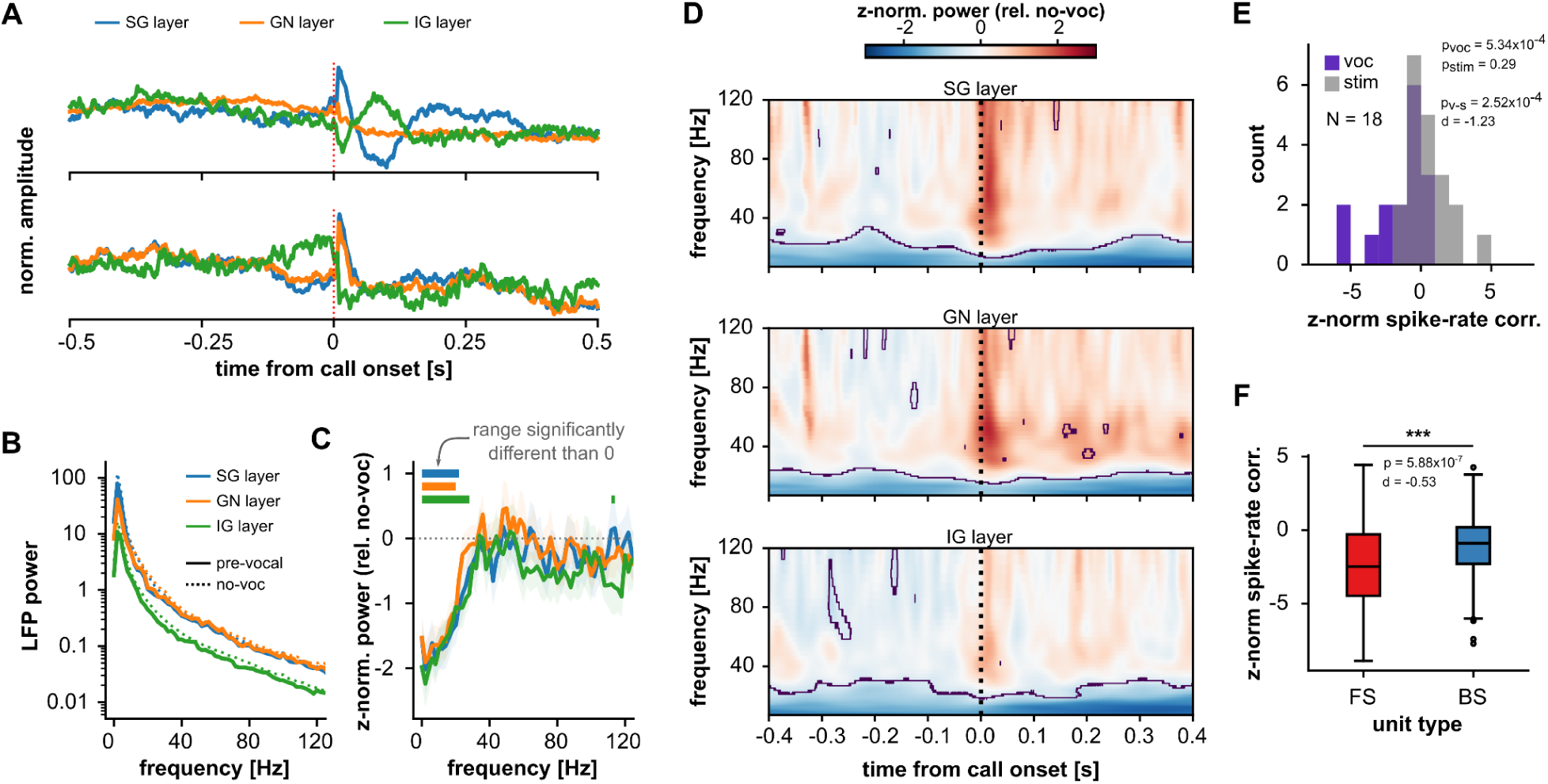
Peri-vocal low-frequency LFP suppression and spike de-correlation in the bat AC. (**A**) Representative peri-vocal LFPs from two different recordings (top and bottom; SG: supragranular LFPs, GN: granular LFPs, IG: infragranular LFPs; see Methods). (**B**) Average pre-vocal LFP power (solid lines) across recordings for the three cortical layers (N_SG_ = 22, N_GN_ = 22, N_IG_ = 17). Dashed lines represent power during no-voc periods. (**C**) z-normalised pre-vocal LFP power relative to no-voc periods across cortical layers. Horizontal lines at the top indicate frequency ranges for which z-normalised power values across recordings are significantly different than 0 (FDR-corrected Wilcoxon signed-rank tests across frequencies, p_corr_ < 0.05). (**D**) z-normalised time-resolved spectrum of peri-vocal LFPs in AC for SG (top), GN (middle), and IG (bottom) layers. Regions with z-normalised power significantly different than 0 across recordings are delimited by contour lines (cluster-based statistics, see Methods). (**E**) Distribution of z-normalised spike firing rate correlations during peri-vocal periods (purple) relative to no-voc activity. The same is shown in gray, but spiking activity was considered during passive listening. Normalised spiking correlations were significantly below zero for peri-vocal periods (Wilcoxon signed-rank test, p = 5.34×10^-4^, N = 18), but not significantly different from 0 during passive listening (p = 0.29). Normalised correlation values were significantly smaller during peri-vocal periods with large effect size (Wilcoxon signed-rank test, p = 2.52×10^-4^, d = −1.23). (**D**) Comparison of normalised spiking correlations across recordings for FS vs. BS units. Both FS and BS units had normalised correlations significantly smaller than 0 (Wilcoxon signed-rank tests, p <= 5.53×10^-14^). Normalised correlation values were significantly smaller for FS units with moderate effect size (Wilcoxon rank-sum test, p = 5.88×10^-7^, d = −0.53).

We further examined LFP power decrease in the time-frequency domain by means of wavelet analyses. Average peri-vocal continuous wavelet transforms (CWT) across recordings are shown in **Fig. 5D**, where CWT power values have been z-normalised to bootstrapped no-voc distributions. Significant power decrease across recordings was observed in all layers for low LFP frequencies (cluster-based statistics; significant regions within contour lines in **Fig. 5D**). Such vocalisation-related power suppression in the LFPs is a sign of low-frequency LFP desynchronisation ^47^. Vocalisation-related LFP desynchronisation was not observed during passive listening (**Fig. S7**), and could not be accounted for by movement alone (**Fig. S5, M-O**).

**Fig 5.**
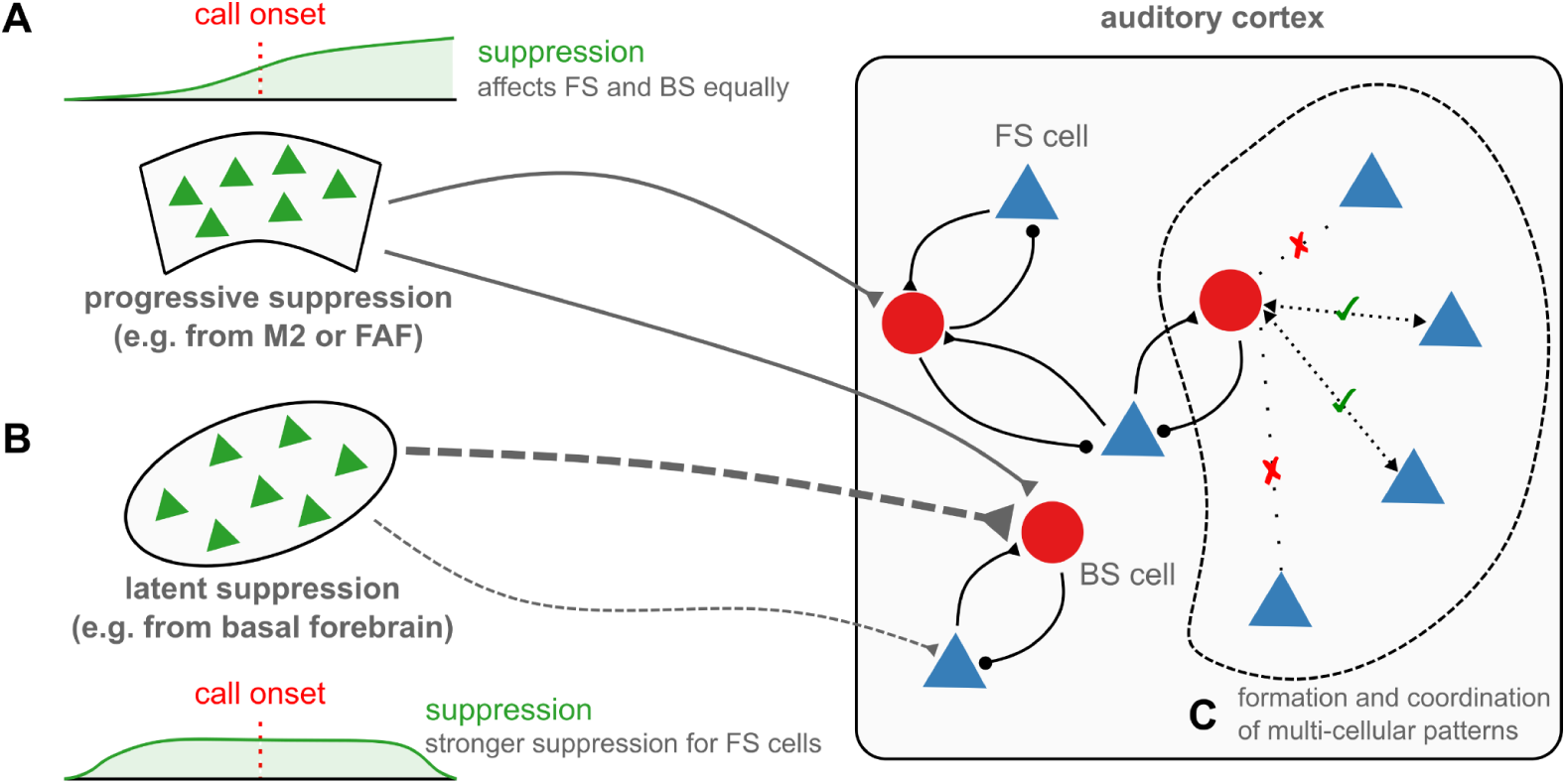
Putative underlying mechanisms for activity patterns of AC fast- and broad-spiking cells during vocalisation. (**A, B**) Two putative sources of vocalisation-related modulation into the auditory cortex, one from modulatory brain centres such as the basal forebrain (**B**), and another one from motor planning or execution regions, such as the secondary motor cortex (M2), the frontal auditory field (FAF), or the laryngeal motor cortex in *C. perspicillata* (**A**). Neuromodulatory regions could cause latent firing rate modulations in the AC, affecting suppressed FS cells most strongly. These modulations could be linked to arousal states. Modulatory signals from motor centres, potentially linked to corollary discharges, could affect firing rates within vocal periods, with suppression gradually increasing over time. These inputs could cause both excitation and suppression in AC. (**C**) FS cells in AC are not only differentially modulated relative to BS neurons but might be instrumental for the formation and coordination of temporally structured firing across multiple neurons.

If there exists a desynchronisation of LFP activity in the AC during vocalisation, we hypothesised that it should be echoed in the neural spiking. We thus measured vocalisation-related spike-rate correlations across neurons for each recording session, and compared them with passive listening and no-voc epochs. Peri-vocal and peri-stimulus correlations were z-normalised using a bootstrapped distribution derived from no-voc neuronal activity. We considered significant deviations from 0 an indication of consistent change in neuronal correlation relative to the no-voc baseline. Z-normalised spike-rate correlations were significantly below 0 (**Fig. 5E**; Wilcoxon signed-rank test, p = 5.34×10^-4^; N = 18), whereas peri-stimulus normalised spiking correlations were not (p = 0.29). Peri-vocal normalised correlations were significantly smaller than during passive listening, with a large effect size (Wilcoxon signed-rank test, p = 2.52×10^-4^, d = −1.23). That is, there was a significant spiking-rate decorrelation in the AC only during vocal production, as predicted by the LFP data. Furthermore, we found evidence indicating that LFP-desynchronisation and spike-rate decorrelations were linearly correlated (Spearman R) over time in a vocalisation trial (**Fig. S7**). In terms of cell type, FS units were more strongly affected by a decrease in spike-rate correlations: FS-related values were lower than those obtained from BS cells, with a moderate effect size (**Fig. 5F**; Wilcoxon rank-sum test, p = 5.88×10^-7^, d = −0.53).

Spike-LFP coherence changes during vocalisation were also cell-type specific (**Fig. S9**). FS cells exhibited no significant change in coherence from no-voc to pre-vocal periods in low-LFP frequencies, as opposed to their BS counterparts. Both cell-types, however, did exhibit an increase in gamma-band spike-LFP synchronisation during pre-vocal periods. We additionally found evidence suggesting that cell-assembly membership could be beneficial for a cell’s spiking in terms of LFP coherence. Altogether, these analyses suggest that the dynamics of FS-cell spiking are more temporally stable than those of BS cells (particularly in pre-vocal periods), in line with their putative role in the formation and maintenance of temporally coordinated activity in AC during vocalisation.

## Discussion

We studied vocalisation-related, cell-type specific neural dynamics in the bat auditory cortex. The results can be summarised as follows: (i) there exist stark differences in the spiking dynamics of putative inhibitory (fast spiking) and excitatory (broad spiking) units related to vocal production, each cell type exhibiting its own modulatory time-course; (ii) novel cellular assemblies and coordinated temporal patterns activate only during vocalisation, and FS units are more likely to participate; (iii) vocal-production induces neural decorrelation, which affects most strongly putative inhibitory cells; and (iv) vocalisation induces spike-LFP synchronisation changes, which differ across neuronal types. Altogether, these results indicate that peri-vocal cortical dynamics exhibit clear cell-type specificity (see **Fig. 5**), and highlight the necessity of considering the richness of cortical heterogeneity for the understanding of vocalisation-related neural activity.

### Cell-type specific cortical modulation of spiking activity during vocal production in AC

In *C. perspicillata*, vocal production induces changes in the firing rate of AC neurons. The modulation of neural activity associated with vocalisation is a well-described phenomenon in animals including mice, human-, and non-human primates ^19–21,51,52^. In marmosets, vocalisation has a predominantly suppressive effect on AC spiking, with excited neuronal subpopulations accounting for less than 40% (occasionally less than 10%) of measured neurons ^25,52,53^. Suppression is not as prevalent in the bat AC: excited cells account for 47% or all significantly modulated units, and excitation and suppression are relatively balanced. Unlike in marmosets, where vocalisation-related excitation occurs after a vocal utterance and appears to be mainly a consequence of sensory reafference ^53^, the majority of excited cells in the bat AC show modulation prior to call onset. Pre-vocal excitation has also been shown for a substantial neuronal population in mice ^20^. Together, observations in bats (this study), rodents, and marmosets suggest interesting differences between vocalisation-related neural modulations across species, and call for additional research to comparatively address this variability.

The AC is generally suppressed during movement and self-produced sounds ^21,23,54–56^. Studies in mice show that movement- and vocalisation-related suppression are largely a consequence of local feedforward inhibition involving parvalbumin-expressing (PV+) interneurons ^21^. This inhibitory circuit is driven by AC-projecting excitatory neurons in M2 which target PV+ cells in all AC layers ^21,57^. M2-AC driven inhibition provides in part a mechanistic foundation for the vocalisation-related suppression reported here. Nevertheless, multiple cells in the bat AC show firing rate modulations that begin before the pre-vocal period (500 ms) examined in this work. While such early modulation could imply that the pre-vocal period is not sufficiently long to capture the onset of the modulatory effect, it could also indicate the presence of complementary modulatory dynamics related to the animal’s behavioural state. Baseline ^58,59^, stimulus ^60^ and task-related ^61^ firing rates not only in AC, but also in other sensory cortices ^62,63^, are altered depending on arousal or engagement ^64^. There exists a cholinergic neuromodulatory circuit wherein basal forebrain neurons, correlated with arousal, target diverse inhibitory and excitatory cell-types in AC evoking both inhibition and excitation ^65^. Although we do not have a read-out of the animals’ state in the current dataset, vocalisation can be a consequence of elevated arousal (e.g. the relationship between vocalisation and Mayer-wave in marmosets: ^66^; see also ^67–69^). Note that accounting for the animal’s movement did not wholly capture the modulatory dynamics observed during vocal production (**Fig. 2, S5**).

We propose that the suppression observed in the current study might have two principal origins. The first (**Fig. 5A**) entails motor related modulatory signals from (pre-) frontal and motor areas, such as the frontal auditory field (a vocalisation-related frontal region anatomically and functionally connected with the AC; ^17,70,71^), the pre-motor cortex (as in mice; ^21^), or the recently described laryngeal motor cortex of *C. perspicillata* ^72^. These signals might be akin to corollary discharges, ultimately suppressing some FS and BS neurons over time and indiscriminately. The second entails a neuromodulatory circuit that alters cortical excitability and firing rates depending on the animal’s arousal state related to vocal production (**Fig. 5B**). When considering suppressive effects, such latent modulation appears to affect FS cells most strongly (**Fig. 2E**, thicker line in **Fig. 5B**). Support for this dual-mechanism hypothesis is found in marmosets, where concurrent and complementary inhibitory dynamics act at different timescales during calling ^73^. Neuromodulatory effects in AC related to arousal and vocalisation might also explain the latent excitation observed in the data ^65^.

In the bat AC, FS units were disproportionately more likely to be modulated than their BS counterparts, and were also significantly more suppressed. Such more selective targeting of auditory cortical interneurons could promote either general inhibition to self-generated sounds, or the refinement of cortical computations (or both) ^29^. The substrate for this might involve neuromodulatory inputs into AC which preferentially modulate inhibitory cell types by means of direct and indirect inhibitory (or excitatory) effects ^65,74^. The FS units described here are more inhibited regardless of whether they show vocalisation-related suppression (**Fig. 2E**) or excitation (**Fig. 2F**), which might ultimately contribute to maintaining overall excitability in cortex, as seen in marmosets ^19^. In bats, an animal that relies on reafferent acoustic cues for navigation, preserving (refined) excitability while engaging and modulating the inhibitory circuitry might be crucial for efficient echolocation-related computations.

### Vocalisation, cell assemblies and temporal spiking patterns

In addition to single-unit dynamics, we characterise the novel activation of cellular assemblies in the AC during vocal production. Cell assemblies are considered a fundamental unit of neural computation and emerge with the coordinated spiking of multiple neurons ^39,40,50,75^. Inhibitory neurons play a critical role in assembly formation, organising the firing of assembly constituents and shaping assembly membership ^76,77^. Given the importance of inhibition and inhibitory cell types in assembly formation and stability, it was predictable that -despite being a minority in our recorded population of neurons- assemblies constituted by at least one FS unit were majoritary. Remarkably, most of the assemblies detected during vocal production were exclusive to peri-vocal periods, and were absent in no-voc or passive listening epochs. Assembly formation during vocalisation likely benefits from the rich peri-vocal firing dynamics of FS and BS cells discussed above.

Analyses of neuronal temporal patterns with SpikeShip ^41^ supported the existence of vocalisation-exclusive, coordinated spiking in the AC. In addition, these analyses reveal that the temporal patterns formed by FS units carry information that is more orthogonal to firing-rate modulations than those of BS cells. This observation is consistent with previous evidence that temporal coding provides information complementary to firing rates in cortical neurons ^78–80^. Our data suggest that the temporal organization of putative inhibitory interneurons is more robust to vocalisation-related modulatory influences. If FS cells are important for the formation and consolidation of cellular assemblies, robustness against vocalisation-related firing rate modulations might confer a higher degree of stability to coordinate firing across multiple neurons. In all, the contrast between FS and BS cells evidenced here indicates that the activity of inhibitory neurons plays a pivotal role in the organisation of AC computations related to vocal production (**Fig. 5C**).

### Neural desynchronisation, decorrelation, and coherence during vocalisation

LFPs provide another measurement of mesoscopic neural activity. Our data reveal a strong suppression of low-frequency LFP power accompanied by firing-rate decorrelations, exclusive to peri-vocal periods, which we interpret as a signature of neural desynchronisation ^47^. Cortical desynchronisation of both LFP and neuronal activities is considered a fingerprint of elevated arousal or attention ^47,64,81^, consistent with the conditions under which an animal could be expected to vocalise ^69^. In such desynchronised states, information processing is enhanced ^82,83^, suggesting that during vocal production the AC is equipped to efficiently deal with the consequences of calling, be it for feedback control or for orchestrating auditory-guided behaviour. The transition to, and maintenance of, desynchronised states is considered to depend heavily on inhibition ^64,84^. Therefore, the fact that FS cells exhibit the highest degree of vocalisation-related decorrelation, together with the observation that they are strongly modulated by vocal production, suggests that AC inhibitory neurons are essential for shaping the circuit dynamics that constitute the hallmarks of vocalisation-related cortical states.

How can neuronal firing-rate decorrelations coexist with the formation and maintenance of temporally coordinated spiking during vocal production? In the case of FS neurons, which were constituents of the majority of vocalisation-related assemblies, multi-neuron spiking patterns showed a high degree of independence from population firing rates (**Fig. 4I**). That is, the firing rate variability across FS cells, arising from firing-rate decorrelations and heterogeneous vocalisation-related modulations (i.e. suppression or excitation throughout the peri-vocal period), only weakly affects the nature of the temporal patterns themselves. A similar argument can be advanced for the case of BS units.

The temporal consistency of AC spiking patterns can be further assessed by referencing spike timings to a secondary signal: the LFP phase. In spite of changes in firing rates and an overall desynchronisation of cortical activity, the temporal relationship between FS units and low/beta LFP phases remained unaltered relative to no-voc periods. The same was not observed for BS cells. These results reinforce the notion that inhibitory neurons in the AC are crucial for establishing and maintaining, with reliability, temporally coordinated spiking during vocal production. Furthermore, spike-LFP phase coherence is thought to be a mechanism for the temporal orchestration of diverse computations in AC, such that the robustness or increase of spike-LFP coherence can be of computational advantage ^44,78,85,86^. Indeed, we report enhanced BS spiking synchronisation with low/beta frequencies, and increased FS- and BS- spiking synchronisation with gamma-band LFPs during pre-vocal periods, as shown for elevated attentional states in previous work ^87–90^. Higher pre-vocal spike-LFP gamma synchronisation in the bat AC could reflect the computational demands of vocalisation imposed on the cortical circuit.

Altogether, this study emphasises fundamental differences in the activities of putative inhibitory versus pyramidal cells in the AC during vocal production, and highlights the particular importance of inhibitory neurons for peri-vocal dynamics. We advance that, rather than considering vocalisation-related neuronal activity as a homogeneous phenomenon, explicitly incorporating the heterogeneity of the cortical circuitry is necessary for the proper understanding of the neural basis of vocal production.

## Methods

### Animal preparation and surgical procedures

This study was conducted on three awake *Carollia perspicillata* bats (two males), obtained from a colony at the Goethe University in Frankfurt am Main. All experimental procedures complied with European regulations and were approved by local relevant authorities (Regierungspräsidium Darmstadt, experimental permit #FR/2007). Animals used in experiments were housed isolated from the main colony, with a reversed light/dark cycle (lights off from 12:00 to 00:00).

Surgical procedures are similar to those described in previous studies from our group ^78,91,92^. Prior to surgery, and for any subsequent handling of wounds, a local anaesthetic was applied subcutaneously over the scalp region (ropivacaine hydrochloride, 2 mg/ml, Fresenius Kabi, Germany). For surgery, animals were anaesthetised with a mixture of ketamine (10 mg*kg−1, Ketavet, Pfizer) and xylazine (38 mg*kg−1, Rompun, Bayer). A rostro-caudal midline incision was cut over the scalp, following which skin and muscle tissue were carefully removed in order to expose the skull. To allow for head-fixation during electrophysiological recordings, a metal rod (ca. 1 cm length, 0.1 cm side) was attached and glued to the bone. The AC was located by means of well-described anatomical landmarks including the sulcus anterior and prominent blood vessel patterns ^17,38,93^). The surface of the AC was exposed by cutting a small hole (ca. 1 mm^2^) using a scalpel blade. Electrophysiological recordings were made mostly in the high frequency fields ^93^ of the left hemisphere.

Post surgery recovery time was no shorter than three days, after which the experimental sessions began. No session was longer than 4 h per day, and only one session per day was done on any animal. Water was given to the animals during a session every 1-1.5 h, and experiments were halted if the animal showed any sign of discomfort (e.g. excessive or violent movement in the holder). Bats were allowed to rest at least three days after four consecutive experimental sessions.

### Electrophysiological, acoustic and video recordings

Electrophysiological recordings were performed acutely on fully awake animals in a sound-proofed and electrically isolated Faraday chamber. Bats were placed on a custom-made holder which was kept at a temperature of approximately 30°C by means of a heating blanket (Harvard homeothermic blanket control unit). Neural activity was recorded with the use of Neuropixels 2.0 electrodes (IMEC, Leuven) in a 4-shank configuration. Channel maps were set such that data were acquired from two banks (closest to the tip) of two shanks, with contacts spanning a total vertical distance of 1425 µm per shank (several hundreds of micrometers longer than the total depth of *C. perspicillata*’s AC; see ^17,94^) and a total of 384 recording sites. The Neuropixels probe was carefully inserted into the brain perpendicular to the cortical surface, and lowered with a piezo manipulator (PM-101, Science products GmbH, Hofheim, Germany) until all contacts were clearly inside the cortical tissue, confirmed both by visual confirmation with binoculars and by online monitoring of signal patterns across channels. The monitoring and online storage of electrophysiological data was done with SpikeGLX (https://billkarsh.github.io/SpikeGLX). Coarse LFP layers (i.e. supragranular, SG; granular, GN; and infragranular, IG) were determined based on three factors: the depth of the Neuropixels probe in the cortical tissue, the activity patterns along the probe (spontaneous and sound-driven), and the well-described reach of auditory cortical layers in *C. perspicillata* based on histological analyses ^17,94^. If the limits of a layer could not be identified with certainty, a recording was not included for the analysis in that particular lamina.

The bat’s vocal behaviour was recorded simultaneously with the neural activity using a microphone (model #4135, Brüel & Kjaer) located 10 cm in front of the animal, amplified with a custom-made amplifier and analog-to-digital converted with a sound card (RME ADI-2-Pro FS; RME Audio, Haimhausen, Germany) with a sampling rate of 352.8 kHz and a depth of 32 bits. Acoustic and electrophysiological recordings were synchronized by means of an external clock signal (generator: DG1022Z, Rigol, China) at 70 Hz, which was fed to the Neuropoixels acquisition system and to one of the input channels of the sound card. Each 3-4 hour recording session consisted of shorter ∼10 minute sub-recordings, because of size limitations inherent to individual *wav* files. Sound recording and presentation (see below) were done in full-duplex mode and were controlled by an in-house developed Matlab (version R2022b; MathWorks, Natick, MA) software.

Animal video recordings were performed with a Basler infrared camera (model: Basler ace acA2000-165um; Basler AG, Ahrensburg, Germany). Acquisition was made at 70 frames-per-second with a resolution of 1080×720 pixels. Each frame capture was triggered by a pulse from the master clock (see above). One video file was produced per ∼10 minute recording. Video acquisition was controlled with the *pylon* software.

### Sound presentation

Natural stimuli were presented to the animals in a probabilistic fashion, such that a sound was played every 2 minutes with a probability of 33% each second (i.e. there was always at least 2 minutes between sound presentations). Two echolocation and two communication calls were used as stimuli. Each stimulus was presented at 70 dB SPL (rms). For each presentation, a stimulus was pseudo-randomly chosen, digital-to-analog converted through the same soundcard used for acoustic recordings, amplified (Rotel power amplifier, model RB-1050) and delivered through a speaker (NeoCD 1.0 Ribbon Tweeter; Fountek Electronics) placed 10 cm away from the animal’s right ear (i.e. contralateral to the measured AC regions). The speaker was calibrated using the same microphone used to record animal vocalisations (see above).

### Data analysis

#### Detection and classification of vocal outputs

Data analyses were performed using custom-written Python (3.10) and Matlab scripts. Vocalisations were automatically detected based on an amplitude threshold algorithm. Prior to detection, the original audio was high-pass filtered at 1 kHz (4th order butterworth filter). For each wav file, a silent period was manually identified from which baseline parameters (mean and standard deviation of the amplitude envelope; µ_*baseline*_ and σ_*baseline*_ respectively) were computed. The envelope of the audio signal was extracted and then z-normalised relative to the baseline period as follows:

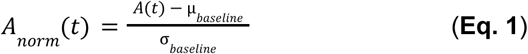

where *A*_*nrom*_(*t*) represents the normalised audio amplitude at time t, and *A*(*t*) the original audio amplitude (high-pass filtered at 1 kHz). A period of audio with a z-amplitude higher than 7 (i.e. 7 standard deviations above the noise floor) and with a duration of at least 0.1 ms was preliminarily deemed a vocalisation. Considering the stereotypical acoustic structure of *C. perspicillata*’s echolocation pulses ^34,37,38^, every tentative vocalisation was classified as “echolocation” or “communication” based on its power spectrum. Preliminary echolocation calls were considered as such if the acoustic event had more than 50% of its spectral energy above 50 kHz, with a duration shorter than 5 ms. If these conditions were not met, an acoustic event was preliminarily considered a communication utterance.

Since we were particularly interested in studying neural patterns during pre-vocal activity, we exclusively considered preliminary calls for which no acoustic event was detected up to 500 ms before call onset (i.e. the pre-vocal period). This subset of preliminary events was then manually curated to refine call classification and to ensure that pre-vocal intervals were free of acoustic contamination, by means of a custom-written Matlab software.

#### Movement quantification and analyses

Movement was quantified as the aggregate difference between consecutive frames in a video. That is, the sum of the differences from all pixels from one frame to the next. To increase SNR relative to the animal’s movement, 1 every 10 frames (i.e. a sampling rate of 7 Hz) were considered. We then applied singular spectrum analysis (SSA) to isolate components that better described the animal’s movement. Only those components were kept for further analysis. Movement quantification was manually curated for each recording session.

We selected silence epochs (no-voc periods, see below) with high-movement or low-movement values. Movement in an epoch was considered the average motion value across all time points in the period. High-movement (HM) epochs were those with the highest 30% motion values; low-movement (LM) epochs were those with the lowest 30% motion values. HM- and LM- no-voc epochs (as many epochs as calls in a recording, 500 bootstrapped repetitions; see below) were chosen from these particular subgroups randomly as described below. Movement during vocalisation periods was calculated as the average movement in the vocalisation trial (i.e. −0.5 to 0.5, re. vocal onset). For comparing movement in no-voc HM or LM epochs, the median across all bootstrapped iterations was considered, and movement values were compared at the level of recordings (FDR-corrected, paired Wilcoxon signed-rank test, significance for p_corr_ < 0.05; **Fig. S5C**). For a given unit we computed the firing rate in each bootstrapped repetition utilising no-voc HM or LM periods. The unit was considered significantly modulated by movement if there were significant differences in firing rate for HM and LM epochs, with at least moderate effect sizes to account for the relatively large number of observations (M = 500 repetitions) yielding significant differences with ease (Wilcoxon rank-sum test, significance for p < 0.05 and |d| ≥ 0.5). Analyses described below, done specifically over the spiking or LFP activity but considering movement-ranked epochs, were performed using the same parameters but considering only the respective neural activity in no-voc HM or LM periods (e.g. LFP power, or SpikeShip analyses).

#### Spiking activity

The handling of raw, spiking and LFP data was done with aid of the *SpikeInterface* (version 0.101.1; ^95^). Spike detection and sorting was performed with Kilosort 4 ^96^. Provisional clusters were manually curated using *phy* (version 2.0b5). These manually curated clusters underwent additional automatic curation with the following parameters (maximum L-ratio = 1, minimum isolation distance = 15; maximum ISI violation percentage = 0.5%; maximum amplitude cutoff = 0.1) computed by the *SortingAnalyzer* module of *SpikeInterface* Spike-sorted units were only considered further if their firing rate was above 1 Hz. To ensure unit stability throughout the analysis period while at the same time maximizing the number of units usable in each recording, per session we kept only consecutive sub-recordings (∼10 minutes in length) in which at least 50% of the units exhibited consistent firing rate across each ∼10-minute period. Under these conditions, the number of consecutive sub-recordings used per session was typically over 10 (i.e. over 1 h 40 min of continuous activity). This restriction of sub-recordings was only done for the analysis of the spiking activity and not for analysing local-field potentials (LFPs; see below). Isolated units were classified into fast-spiking (FS) and broad-spiking (BS) groups based on two metrics: peak-to-valley duration and repolarization slope, computed with *SpikeInterface*’s *SortingAnalyzer* module. We verified unit separability in terms of peak-to-valley duration by means of a Hartigan’s dip test (uncalibrated p = 0.053; calibrated p = 0.0001; based on the repository for ^97^) and defined a cutoff at 460 μs for this metric (i.e. units below that cutoff were considered FS; above, BS). We additionally verified unit classification by means of a k-means clustering algorithm (2 clusters) with excluded outliers wherein both peak-to-valley and repolarization slope were considered, and observed good separation of spike waveform properties. The outcome of the clustering and the peak-to-valley cutoff were both considered to determine whether a unit was FS, BS, or not-classified (NC). These analyses are summarised in **Fig. S2**.

For each unit we calculated instantaneous firing rates during peri-vocal activity (±500 ms around call onset) using *zetapy*’ instantaneous firing rate (IFR) as it avoids the need for binning the spiking data (https://github.com/JorritMontijn/zetapy, see also ^98^). For comparability across neurons, firing rates were interpolated to 5 ms, given that IFR time-vectors differ depending on the temporal dynamics of the spiking rate for each cell. In addition to peri-vocal IFRs, we obtained IFRs from pseudo-randomly selected no-voc periods in which the animals emitted no calls and in which there was no other form of acoustic contamination. The number and duration of no-voc periods matched the number and duration of vocalisation trials. To avoid sampling biases, no-voc periods were selected M = 500 times (yielding 500 no-voc IFR traces), from which we derived the mean and standard deviation of baseline spiking rates.

We determined whether a unit’s firing rate was significantly modulated during peri-vocal periods based on cluster-based statistics ^100^ combined with a jackknife approach. First, we z-normalised (see **Eq. 1**) a unit’s peri-vocal IFR using as baseline M - 1 no-voc periods. Initial candidate clusters were periods of at least 20 ms for which the absolute z-IFR was above 1.96 (equivalent to p = 0.05). The cluster-mass for each cluster was defined as the sum of the z-normalised values of IFR for each time point in the cluster, and compared to a bootstrapped distribution of cluster-masses obtained using a jackknife procedure as follows. For each no-voc iteration *m*, we normalised its IFR using the remaining M - 1 no-voc periods as described above. We then determined the cluster-mass of the significant clusters in that particular bootstrapping iteration and chose the largest. Across all bootstrap iterations, 500 cluster-masses were computed which were used as a null distribution against which the original cluster-masses were compared. A cluster observed during peri-vocal periods was deemed significant if its cluster-mass value was in the 99^th^ percentile of the bootstrap null distribution. Units with significant clusters were considered to be significantly modulated in firing rate during vocalisation trials. If the cluster-masses were predominantly negative (i.e. peri-vocal IFR below spontaneous firing rate) the unit was considered to be suppressed, and if they were predominantly positive the unit was considered to be excited. Units which showed two types of modulation were atypical, but for those which did we considered the sum across all cluster-masses to be the defining factor in the classification.

#### Local-field potentials

LFP signals were computed by bandpass filtering (0.01 – 300 Hz) and downsampling (to 5000 Hz) the raw data across electrodes by means of *SpikeInterface*’s pre-processing tools. For a particular recording, layer-specific LFPs were computed as the mean across all channels that belonged to a each given layer (i.e. SG, GN, or IG; see above). Pre-vocal LFP power was calculated using Welch’s method with a window length of 500 ms. Time-frequency resolved spectra were computed by means of wavelet transforms (package *PyWavelets*, version 1.8.0) using complex Morlet wavelets with centre frequency and bandwidth of 0.6 and 1.7, respectively, and 120 evenly log-spaced scales between 1 and 1200. The trial-related (i.e. pre- or peri-vocal) power for each recording was defined as the mean power across all calls considered for that particular session.

Vocalisation-related LFP power (computed by Welch’s method or wavelet transforms) per recording was z-normalised (see **Eq. 1**) using as baseline a bootstrapped spectrum. This baseline spectrum was obtained from pseudo-randomly selected no-voc periods (see above) matching in number and duration the vocalisation-related periods. To avoid sampling biases, the pseudo-random selection of no-voc periods was repeated 250 times. The mean and standard deviation of the baseline power was obtained across all bootstrap repetitions. Spectral calculation parameters for no-voc periods were exactly the same as those performed for vocalisation trials.

LFP spectra associated to no-voc periods were used to determine whether there were power modulations in the field potentials during peri-vocal periods. In the case of pre-vocal power spectra (obtained with Welch’s method), we z-normalised the spectrum for each recording session and layer (SG, GN, IG) using the bootstrapped no-voc spectra (M = 250 repetitions). We reasoned that any consistent power change from baseline across all recordings would result in significant differences from 0 for the z-normalised power in a given frequency range (FDR-corrected Wilcoxon signed-rank tests for each frequency in the spectrum, significance if p_corr_ < 0.05). In the case of the wavelet transforms, we determined significant time-frequency regions by means of cluster-based statistics similar to those described for the spiking activity. Spectral values were z-normalised relative to the baseline distribution of no-voc periods (M = 250 repetitions, in this case we used M - 1 no-voc repetitions) and a z-scored wavelet spectrum was obtained for each recording. As above, we considered any significant deviation from 0 in the z-score spectra as evidence for consistent differences between peri-vocal and no-voc periods across recordings. Thus, we found preliminary regions (clusters) in the time-frequency domain for which the z-scored power for contiguous points was significantly different than 0 (Wilcoxon signed-rank tests, p < 0.05) by means of a flood-fill algorithm. The cluster-mass of each region was the sum of the t-statistic at each time-frequency point. Then, for each no-voc repetition *m*, we normalised its spectrum with the remaining M - 1 no-voc repetitions and obtained the maximum cluster-mass across bootstrap iterations. This yielded 250 cluster-mass values which were used as a null distribution against which the peri-vocal cluster-masses were compared. Clusters with cluster-masses more extreme than the 95^th^ percentile of the null distribution were considered to be significant.

#### IFR comparison between FS and BS cells

Peri-vocal firing rates between FS and BS units were compared to determine differences in their modulation patterns over a vocalisation trial by means of cluster-based statistics. For example, to compare between excited FS and BS units (note that the same applied for suppressed units), we determined clusters in which the IFRs significantly differed (two-sample independent t-tests over time, significance for p < 0.05) and computed their cluster-mass as the sum of the t-statistic for each time point. We then pseudo-randomly switched labels between FS and BS units (respecting the original group sizes) a total of 1000 times, and computed a null distribution of maximum cluster-masses that was used as a baseline for comparing peri-vocal cluster-masses (see above). Cluster-mass values more extreme than the 95^th^ percentile of this null distribution were considered significant.

#### Spike-rate correlations

Spiking correlations were computed for each recording individually using the *Elephant* package (version 1.1.1; ^101^), using a bin size of 15 ms. Correlations were computed for all unit pairs in a given recording during peri-vocal, peri-stimulus, and no-voc periods (in the latter, as above, we repeated the calculations 500 times over pseudo-randomly selected segments). The average correlation across all neuron pairs was then taken as the recording’s correlation. Spike-rate correlations for FS and BS units were calculated separately as the mean correlation of every FS or BS unit with every other unit in a particular recording. Z-scored correlation values were obtained by normalising peri-vocal or peri-stimulus correlation values relative to those obtained for no-voc periods (M = 500 repetitions).

A time-resolved spike-rate correlation was also computed for each recording, using a sliding window of 200 ms over the peri-vocal period with a step of 20 ms. In order to determine whether the time-course of the spike-rate correlation covaried with the LFP desynchronization (i.e. a suppression of LFP power in a certain band), we calculated the Spearman correlation between the band-limited instantaneous energy of the LFP for each layer and the time-resolved spike-rate correlation. Band limited LFP energy (in frequencies 2-20 Hz, and 60-90 Hz) was computed as the absolute value of the Hilbert transform of the band-pass filtered LFP (4^th^ order Butterworth filter). LFPs were bandpass filtered using the filtering tools of the *neurodsp* package (version 2.3.0; ^102^).

#### Pairwise phase consistency

Spike LFP synchronization was calculated with the pairwise-phase consistency (PPC) metric ^48^. The PPC is equivalent to the population statistic of the phase-locking value, but it allows computing spike-LFP synchronization in a way that is bias-free relative to sample size (i.e. number of spikes) and firing rates. The PPC was computed for every unit relative to LFP phases in three frequency bands: 4-10 Hz (low frequency), 12-20 Hz (beta), and 60-90 Hz (gamma). The LFP associated to a unit (i.e. from the same electrode) was filtered using *SpikeInterface*’s pre-processing module (see above) in the frequency band of interest, and its instantaneous phase was computed by means of a Hilbert transform. The spike phases were then obtained as the phase of the LFP at each spike time. Only units that exhibited significant circular non-uniformity were considered for PPC comparisons (Rayleigh test of uniformity, significance for p < 0.05; in the package *astropy*, version 6.1.7; ^103^). In addition, considering that the sample estimator of the metric can yield negative values ^48^, we only kept units whose PPC values were strictly non-negative.

Furthermore, a unit’s PPC was only calculated for a given frequency band if we found evidence to support the presence of elevated power in that band. That is, if the power for those frequencies was significantly higher than what is to be expected from fitting a 1/f function on the power spectrum of the LFP from the unit’s channel. Based on the methodology described in ^49^, we found the 1/f fit to the LFP measured at the unit’s channel independently for each of the sub-recordings in a session (typically numbering 12-16), using a spectral parametrization model ^104^. The LFP spectrum was normalised relative to the 1/f fitted spectrum, and we then tested for each frequency whether the normalised spectrum was consistently above 0 across subrecordings (One-sample t-test for each frequency, significant if t-statistic > 2) for at least 2 consecutive frequency points. We took cases of significance within the frequency band of interest as evidence for the presence of consistent, elevated power in the LFP for that given channel.

In addition to pre-vocal PPC, we calculated for each unit a bootstrapped distribution of no-voc related phase-consistency values, following the same randomization procedures described above (M = 250 repetitions). Vocalisation-related spike-LFP synchrony was then z-normalised to the no-voc distribution. We considered significant deviations from 0 in a population of z-normalised PPC values (Wilcoxon signed-rank tests, significance for p < 0.05) as evidence of consistent change in spike-LFP synchronization during pre-vocal periods relative to spontaneous activity.

#### Cell assemblies and spike patterns

Cell assemblies were computed with *Elephant*’s cell assembly detection module, which capitalizes on the analysis in ^40^. This approach is designed to detect cell assemblies with arbitrary organization at various timescales. For detection, we used a bin size of 15 ms and a maximum time lag of 75 ms. Assemblies were detected using all units on a recording-per-recording basis during peri-vocal, peri-stimulus, and no-voc periods. No-voc procedures resembled closely those described above: we chose an equal number of no-voc trials as of vocalisation trials, and used those to identify assemblies (M = 250 repetitions).

Assemblies that did not appear in any no-voc repetition or any peri-stimulus trial were considered unique to vocalisation periods. Only assemblies with activation times in pre-vocal periods were considered (206 of 207 assemblies).

The synchronization of cell assemblies with the LFP was computed in a similar way to that of spike-LFP. As an LFP signal for the assembly, we used the average LFP across the channels from the assembly’s constituent units. Instead of spike times, we used assembly occurrence times and derived assembly occurrence phases from the phase time series of the LFP in low, beta and gamma frequencies. These phases were used to calculate the assembly’s PPC. Significant synchronization to the LFP was assessed with a Rayleigh test (p < 0.05).

The presence of spike temporal patterns was further assessed by means of *SpikeShip*, a firing-rate independent, binless, and unsupervised method that computes dissimilarity across the firing patterns of multiple units ^41,42^. In short, SpikeShip computes how similar all the relative spike-timing relationships are between any two time epochs. Because the metric benefits from a large number of neurons (i.e. > 100; see ^42^), the spiking activity of all units was pooled as follows. First, spiking was pooled across units utilizing 30 of their vocalisation trials (units with less vocalisation trials were not included; 30 trials were pseudo-randomly chosen in cases where more trials were available), with spike times centred at the vocalisation time in a window spanning ± 500 ms. We then defined 30 pre-vocal and 30 post-vocal epochs from each vocalisation trial. Matching the trial selection for vocalisation, we pseudo-randomly chose no-voc epochs, choosing two groups of 30 no-voc periods with a duration of 500 ms. SpikeShip dissimilarity was computed for the spiking train of pooled units across these epochs. We repeated the above procedure a total of M = 250 times, to minimize sampling biases caused by trial selection. The mean dissimilarity matrix across repetitions was directly used to illustrate pattern formation by means of a t-distributed stochastic neighbour embedding (t-SNE). Pre-voc, post-voc and no-voc dissimilarities were obtained as the mean dissimilarity in the dissimilarity matrix corresponding to the period of interest, thus yielding as many dissimilarity values as repetitions (250) for each epoch type. These were compared using FDR-corrected rank-sum tests (significance when p_corr_ < 0.05). Note that by selecting units across multiple recordings and pooling them into a pseudo-simultaneous population with spikes centred on call onset, the temporal spiking patterns dynamics being measured are dominated by those locked to vocalisation utterances.

A similar analysis was performed separately for FS and BS units. In addition to the dissimilarity matrix obtained from temporal spiking patterns, we computed a dissimilarity matrix based on firing rate across epochs and repetitions (see ^42^). As a measure of the relationship between temporal spiking patterns and firing rate variations, we calculated the correlation of both dissimilarity matrices (yielding as many correlation values as repetitions, M = 250). FS-related correlation values were statistically compared (Wilcoxon rank-sum test, significance for p < 0.05) to the BS ones after undergoing Fischer’s z-transformation.

### Statistical procedures

Statistical comparisons were typically performed with paired or unpaired Wilcoxon signed-rank or rank-sum tests, respectively. All statistics, unless otherwise noted, were corrected for multiple comparisons with the False Discovery Rate (FDR) using the Benjamini and Hochberg procedure ^105^. The threshold of statistical significance was an alpha of 0.05. As an effect size metric we used Cohen’s d:

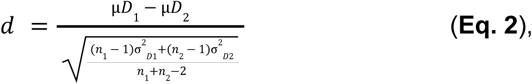

where D1 and D2 represent the two distributions being compared, µ represents their mean, σ^2^ represents their variance, and n_1,2_ their sample sizes. Following ^106^, effect sizes were considered small for |d| < 0.5, moderate for 0.5 ≤ |d| ≤ 8, and large for |d| > 0.8.

Statistical analyses for time series (e.g. LFPs or firing rates), Welch frequency spectra or time-resolved spectra (wavelet transforms) often involved bootstrapping and cluster-based analyses ^100^. These are detailed in their corresponding Methods subsections. When comparing two populations directly, as in the case of FS vs BS firing rates, two-sample independent t-tests were performed for each time point, given that the t-statistic can be linearly summed to obtain cluster-masses ^100^.

## Acknowledgments

This work was supported by the DFG.

## Conflict of interests

The authors declare no financial or non-financial conflict of interest.

## Data availability

The data and code used in this study are available from the authors upon reasonable request.

## Author contributions

F.G.R. and J.C.H. conceived and designed the research. F.G.R. collected the data, analyzed it, produced figures, and wrote the original manuscript. F.G.R., B.S.G, D.P, and J.C.H. discussed analyses and results, interpreted the data, and reviewed figures and text.

## Supplementary materials

**Fig. S1.**
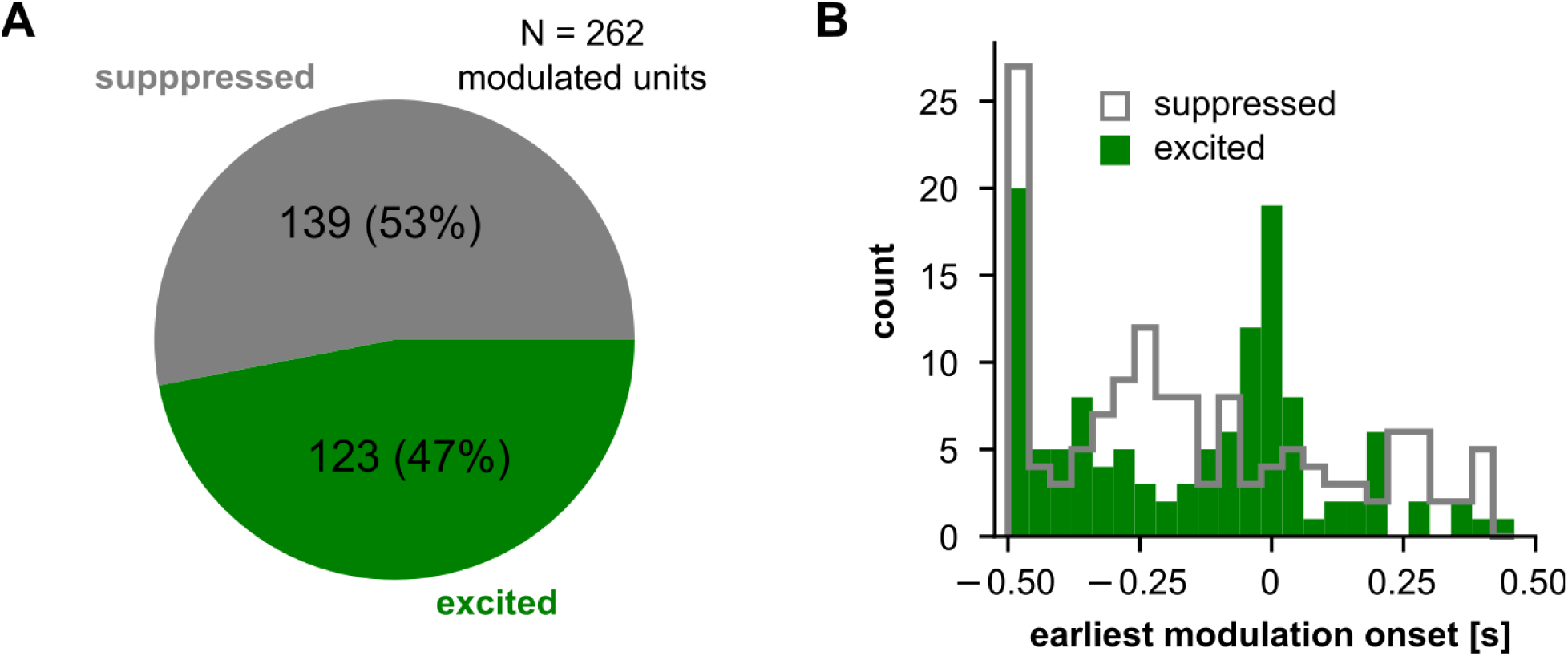
Distribution of vocalisation-related modulations in AC. (**A**) Proportion of suppressed and excited units in the vocalisation-related populations. (**B**) Distribution of earliest modulation onset (the onset of the first significantly modulated cluster across units) for suppression and excitation.

**Fig. S2.**
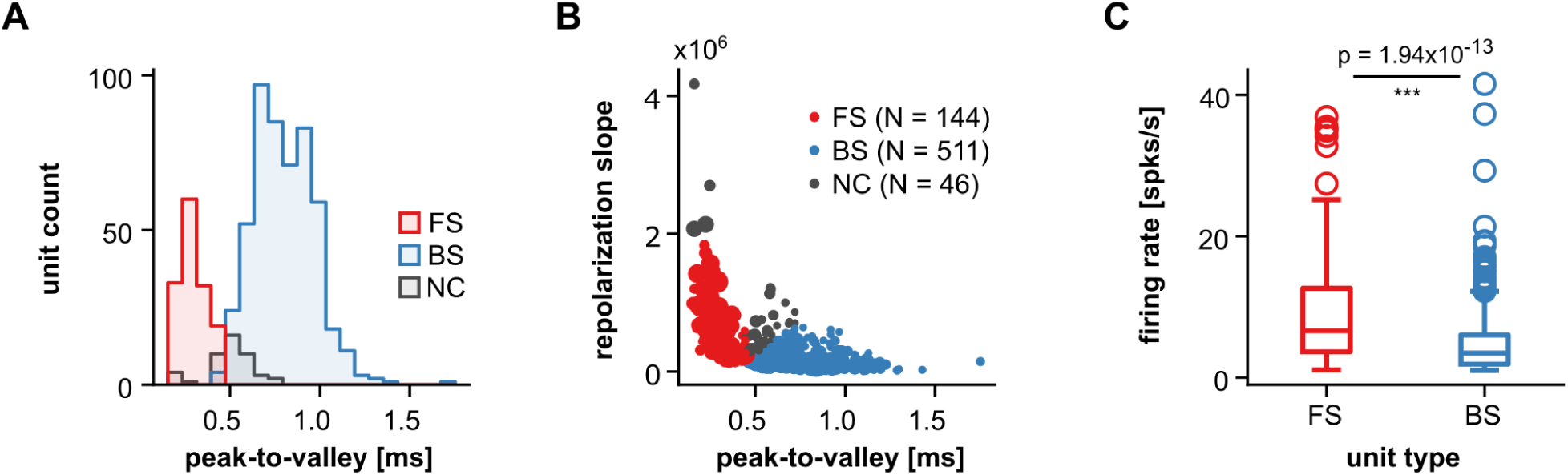
Spiking properties of FS and BS units. (**A**) Distribution of peak-to-valley durations across neuron types. (**B**) Peak-to-valley and repolarisation slopes for all units (N = 701). Each circle represents a unit, with size depending on its firing rate. A k-mean clustering using these two variables, together with a 460 µs threshold for peak-to-valley duration (as in **A**) were used for unit classification (see Methods). (**C**) The firing rates of FS units were significantly higher than those of their BS counterparts (Wilcoxon rank-sum test, p = 1.94×10^-13^).

**Fig. S3.**
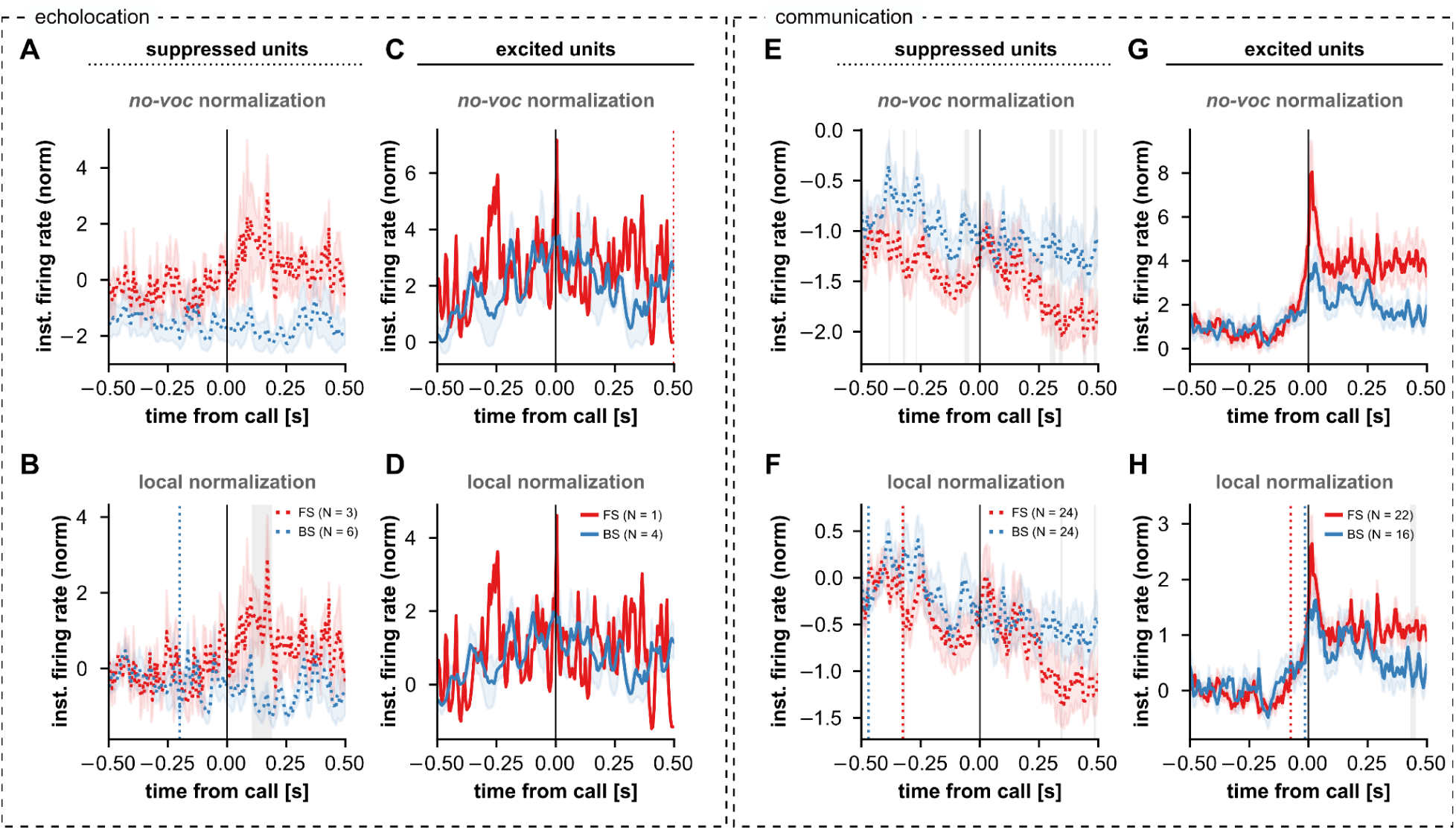
Call-type specific, peri-vocal firing patterns of FS- and BS units. (**A-D**) Follows **Fig. 3A-D**, but showing peri-vocal firing patterns of significantly modulated FS and BS units when animals emitted echolocation calls. (**E-H**) Same as **A-D**, but peri-vocal spiking patterns were considered only when animals vocalised communication calls. The sample sizes were not sufficient to draw conclusions from analysing peri-vocal activity in a call-specific manner, even when relaxing the minimum number of trials necessary for a unit to be considered (here: at least 10 vocalisations; in the main analysis: at least 15). For example, in the echolocation case only 1 excited FS cell was found (panels **C, D**). The reason is the sparseness of echolocation calling in our sessions. Indeed, most echolocation utterances were found in a small number of recordings with relatively low neuronal yield. In other recordings, echolocation utterances contributed to the trial pool, but were not enough to be analysed on their own on a neuron-per-neuron basis. Therefore, and in order to maximise the number of vocalisation trials and the number of vocalisation-modulated units considered, echolocation and communication calls were both considered as “vocalisations” in the main text.

**Fig. S4.**
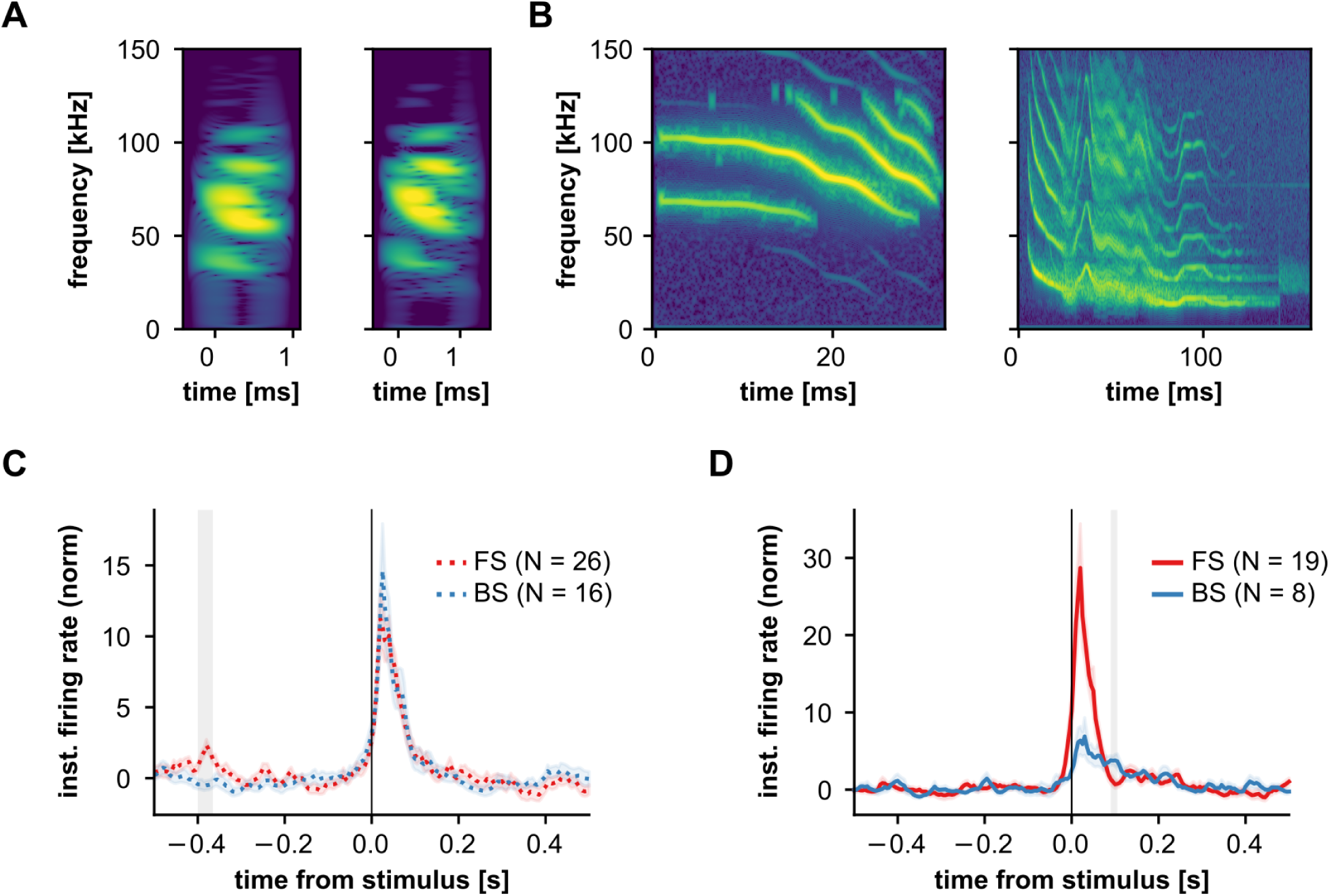
Spiking activity of vocalisation-modulated units in response to acoustic stimulation with natural calls. (**A, B**) Natural stimuli used for acoustic stimulation: two echolocation (**A**) and two communication (**B**) calls. (**C**) IFRs of significantly stimulus-responsive (cluster-based statistics) FS (red; N = 26) and BS (blue) units, normalised to a no-voc baseline. These units were in addition significantly suppressed during vocal production. Firing rates were not significantly different throughout the stimulation period (cluster-based statistics), except for a significant time segment shown as a gray area at about −400 ms. Without stimulation at those time-points, the interpretation for this significant cluster is unclear. (**D**) Same as **C**, but considering vocalisation-related excited FS and BS units. Apparent differences between FS and BS units immediately after vocalisation onset were not significant (cluster-based statistics), possibly due to firing rate variability and relatively small sample sizes (FS, N = 19; BS, N = 8). In both **C** and **D**, significant stimulus responsiveness was determined using cluster-based statistics similar to those used to determine significant vocalisation-related modulations. Differences for the IFRs across cell types were statistically assessed using the same paradigm as that of **Fig. 3**.

**Fig S5.**
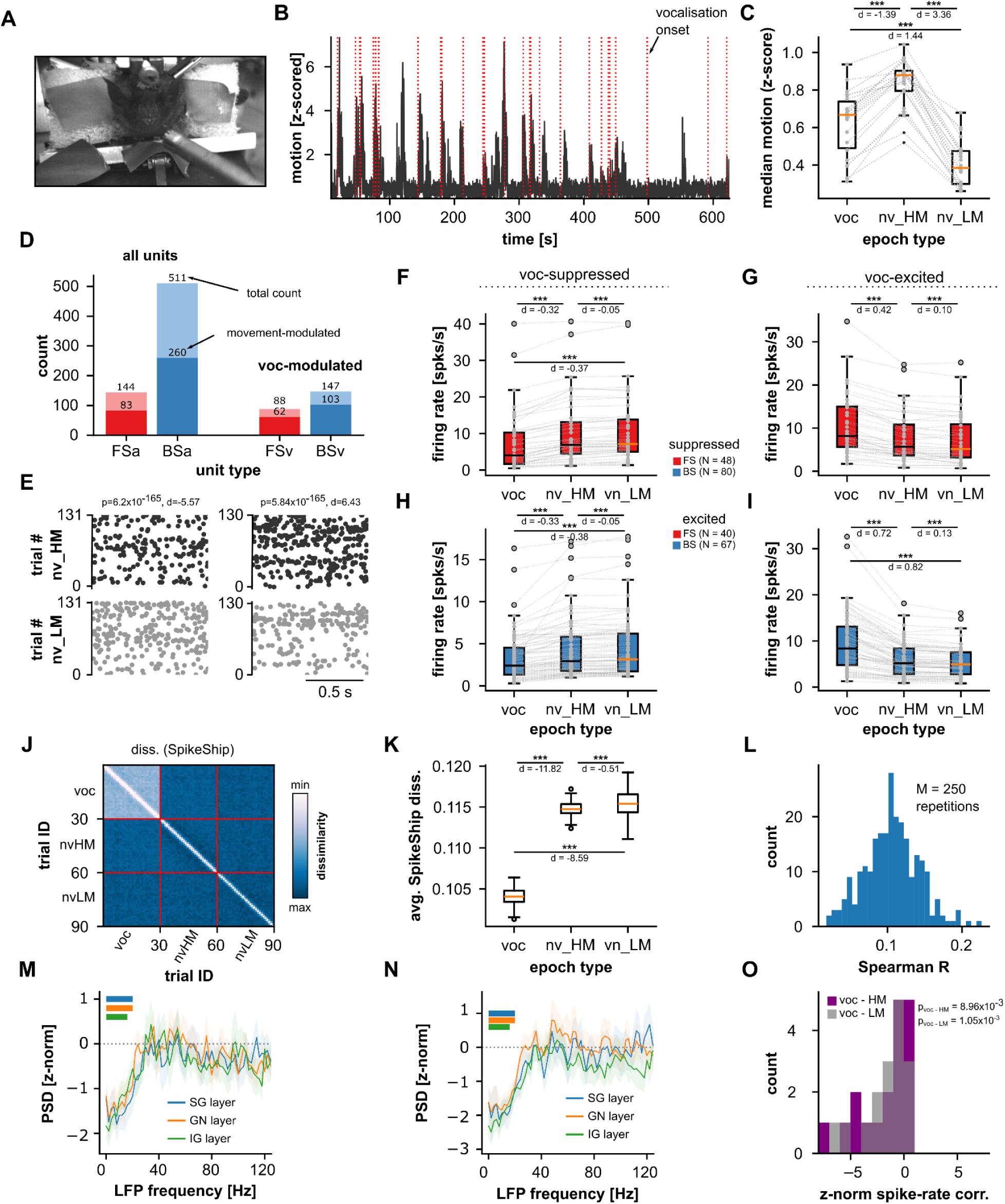
Movement alone does not explain vocalisation-related neural dynamics. (**A**) A still-frame corresponding to one session’s video recording. (**B**) Motion quantification for a 10-minute sub-session in a representative recording. Vocalisation onsets are marked with red dashed lines. Vocalisation is paired with movement, but movement also occurs at times when no calls are uttered. (**C**) Comparison of motion in three epoch types, across all recording sessions (N = 22): vocalisation (voc), no-voc high-motion (nv_HM; i.e. no-voc epochs with the highest 30th percentile of quantified motion), and no-voc low-motion (nv_LM; no-voc epochs with the lowest 30th percentile of quantified motion). Movement during vocalisation was significantly higher (with large effect size) than during nv_LM periods (Wilcoxon signed-rank test, p = 9.54×10^-7^, d = 1.44). Note, however, that movement during nv_HM epochs was the highest of all epoch types (also than vocalisation), with large effect sizes (p ≤ 7.1×10^-7^, |d| ≥ 1.39). (**D**) Number of units in the AC, modulated by the animal’s movement, according to cell type (FS, red; BS, blue). *Left*: considering all of the units; *right*: considering only units that were significantly modulated by vocalisation. Total and relative numbers are indicated in the figure. 58% of all FS units (and 70% of vocalisation-modulated FS units) were modulated by movement. 50% of all BS units (and 70% of vocalisation-modulated BS units) were modulated by movement. A unit was considered modulated by movement if the firing rate in nv_HM epochs was different than in nv_LM ones (Wilcoxon rank-sum test, p < 0.05), with at least moderate effect sizes (|d| >= 0.5). (**E**) *Left*: Raster plots for an example movement-modulated unit (top, nv_HM, dark spikes; bottom: nv_LM, gray spikes). One repetition of nv_HM and nv_LM bootstrap iteration is shown (see Method). Note that this unit’s firing rate decreased in nv_HM epochs (indicated by negative d, shown on top). *Right*: A similar representation, but for an exemplary unit whose firing rate increased during nv_HM trials relative to nv_LM ones. (**F, H**) Firing rates for vocalisation-suppressed cells during voc, nv_HM, and nv_LM epochs, shown for FS (**F**) and BS (**H**) units. (**G, I**) Same as in **F,H**, but vocalisation-excited units are shown. These units are the same used in **Fig. 3** of the main text. Vocalisation-related modulations were always significantly stronger (either for suppression or excitation) than in either high- or low-movement no-voc epochs (Wilcoxon signed-rank test, p ≤ 2.45×10^-9^, |d| ≥ 0.32). Note that this occurs even when movement in nv_HM was stronger than in vocalisation epochs. Thus, firing rate modulations observed in our dataset are not wholly accounted for by the animal’s motion. (**J**) Median SpikeShip dissimilarity matrix (see **Fig. 4**) considering epochs of vocalisation, nv_HM, and nv_LM. (**K**) Spike temporal patterns were significantly more stable during vocalisation epochs than during either high- or low- movement epochs, with large effect sizes (Wilcoxon rank-sum test, p ≤ 3.33×10^-83^, d < −8.6). (**L**) Distribution of the correlation between SpikeShip and and firing rate dissimilarity matrices, across repetitions (M = 250). Relatively low correlation values indicate independence between the temporal patterns and the neuronal firing rates. Altogether, panels **J-L** indicate that the neuronal temporal structure observed during vocalisation (**Fig. 4**) cannot be fully accounted for by movement. (**M**) Z-normalised pre-vocal LFP power relative to hv_HM periods across cortical layers. Horizontal lines at the top with colours matching layer identifying colours, indicate frequency ranges for which z-normalised power values across recordings are significantly different than 0 (FDR-corrected Wilcoxon signed-rank tests across frequencies, p_corr_ < 0.05). (**N**) Same as in **M**, but pre-vocal LFP power was normalised to nv_LM epochs. (**O**) Distribution of z-normalised spike firing rate correlations during peri-vocal epochs, relative to nv_HM (purple) or nv_LM (gray). Normalised spiking correlations were significantly below zero in both cases (Wilcoxon signed-rank test, p ≤ 8.96×10^-3^, N = 18 recordings). Panels **M-O** indicate that the neural decorrelation, observed in the LFP and the spiking activity (**Fig. 5**), cannot be fully explained by the animal’s movement.

**Fig. S6.**
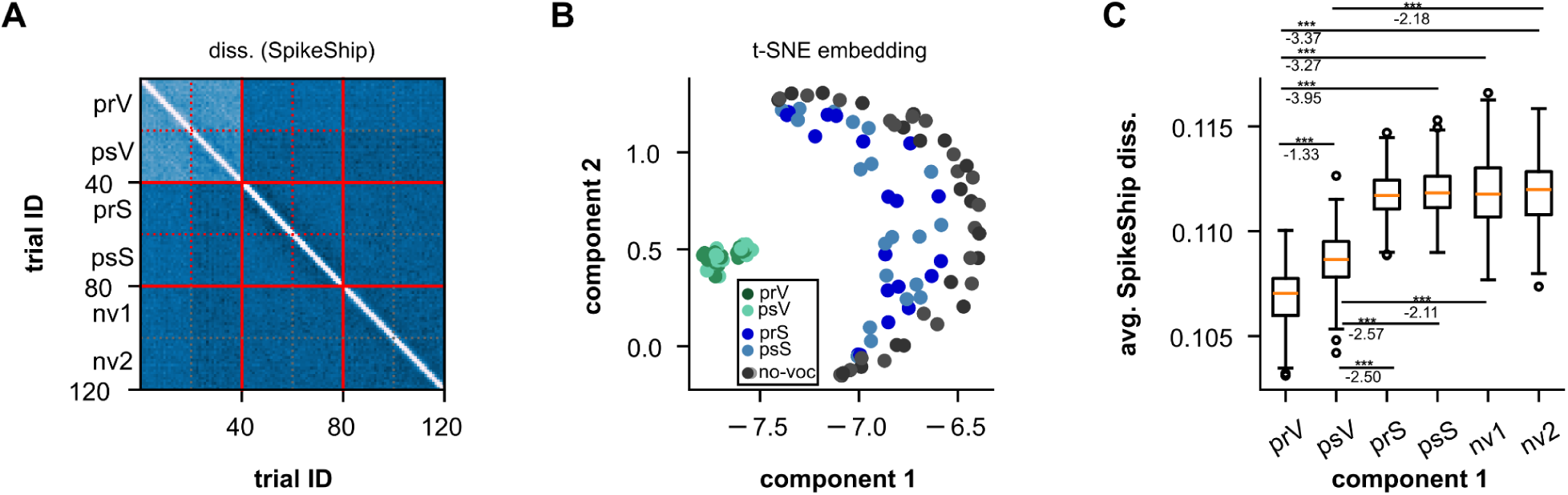
SpikeShip pattern analysis including epochs of acoustic stimulation with natural stimuli. (**A**) Average SpikeShip dissimilarity matrix across repetitions (M = 250), for epochs that include pre-vocal, post-vocal, pre-stimulus, post-stimulus, and no-voc trial types. The SpikeShip analysis is similar to the one of **Fig. 4** in the main text. However, 20 epochs (instead of 30) of each type were selected, and units were only included if they had sufficient vocalisation and passive listening trials (i.e. at least 20). Because of these requisites, only a subset of 282 units were used (in the main analysis, 417 were used). (**B**) 2-dimensional t-SNE embedding obtained from the dissimilarity matrix in **A**. (**C**) Average SpikeShip dissimilarity across trial types. Values for pre-vocal and post-vocal epochs were significantly smaller than for pre-/post-stimulus and no-voc trials with large effect sizes (FDR-Corrected Wilcoxon rank-sum tests, p_corr_ < 9.80×10^-63^; d < −2.11). SpikeShip dissimilarities were also different for pre-vocal and post-vocal trials (p_corr_ < 3.42×10^-37^; d < −1.33). Altogether, these results support that spike temporal patterns formed during vocal production are unique to this behavioural state.

**Fig. S7.**
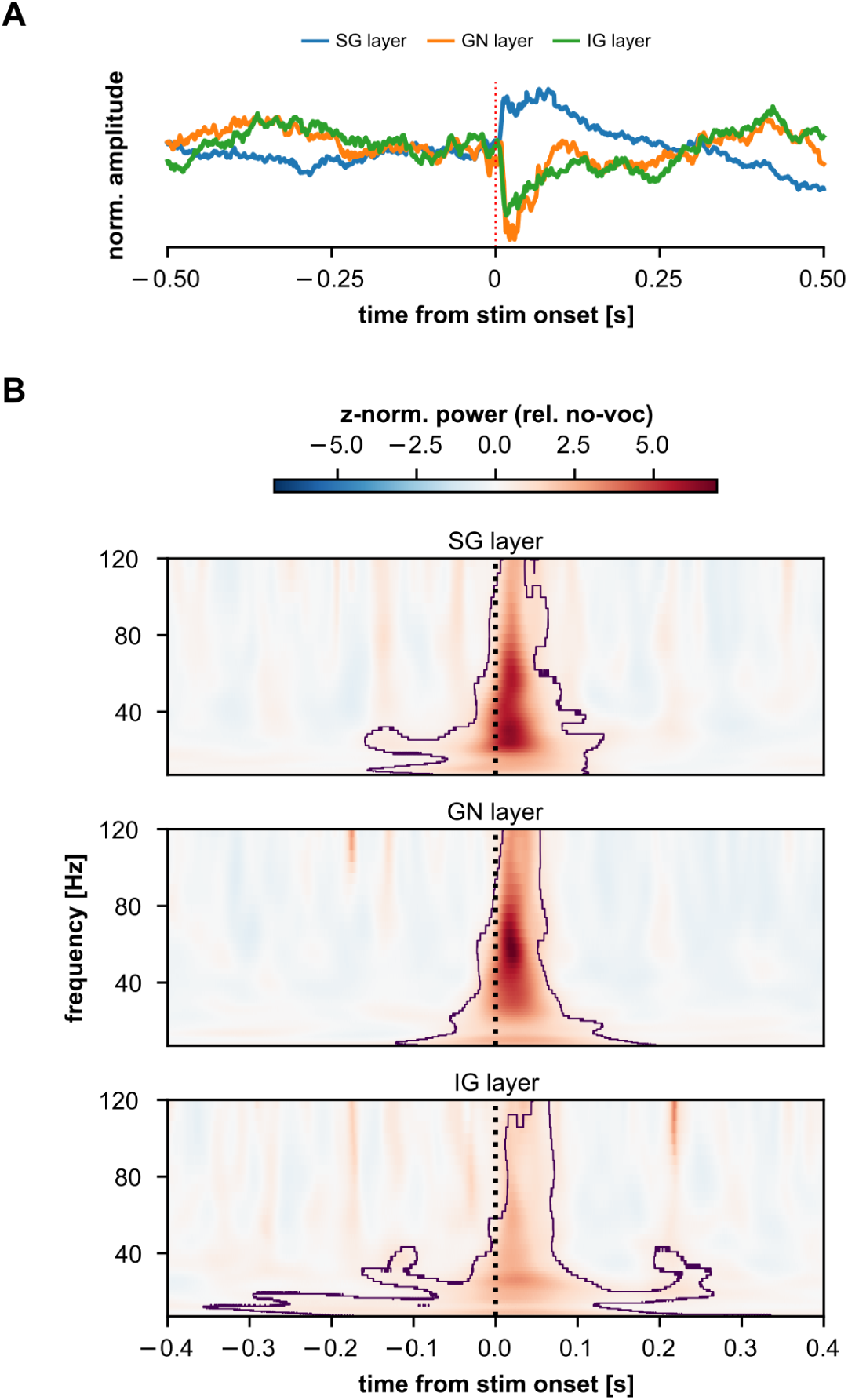
LFP response to acoustic stimulation. (**A**) Stimulus-evoked LFPs from a representative recording session (SG: supragranular LFPs, GN: granular LFPs, IG: infragranular LFPs). (**B**) Stimulus-evoked time-frequency resolved LFP spectra across cortical layers, averaged across recording sessions. The treatment to these spectra is the same as that shown in **Fig. 5D**: they were z-normalised to a no-voc baseline. Significant deviations from 0 across recordings were interpreted as a consistent trend in our dataset. Contour lines in the figure delineate significant clusters (see Methods). A clear evoked-related pattern indicates typical primary-like auditory responses to the natural stimuli. No evidence for low-frequency LFP desynchronization is visible during passive listening (cf. **Fig. 5**).

**Fig S8.**
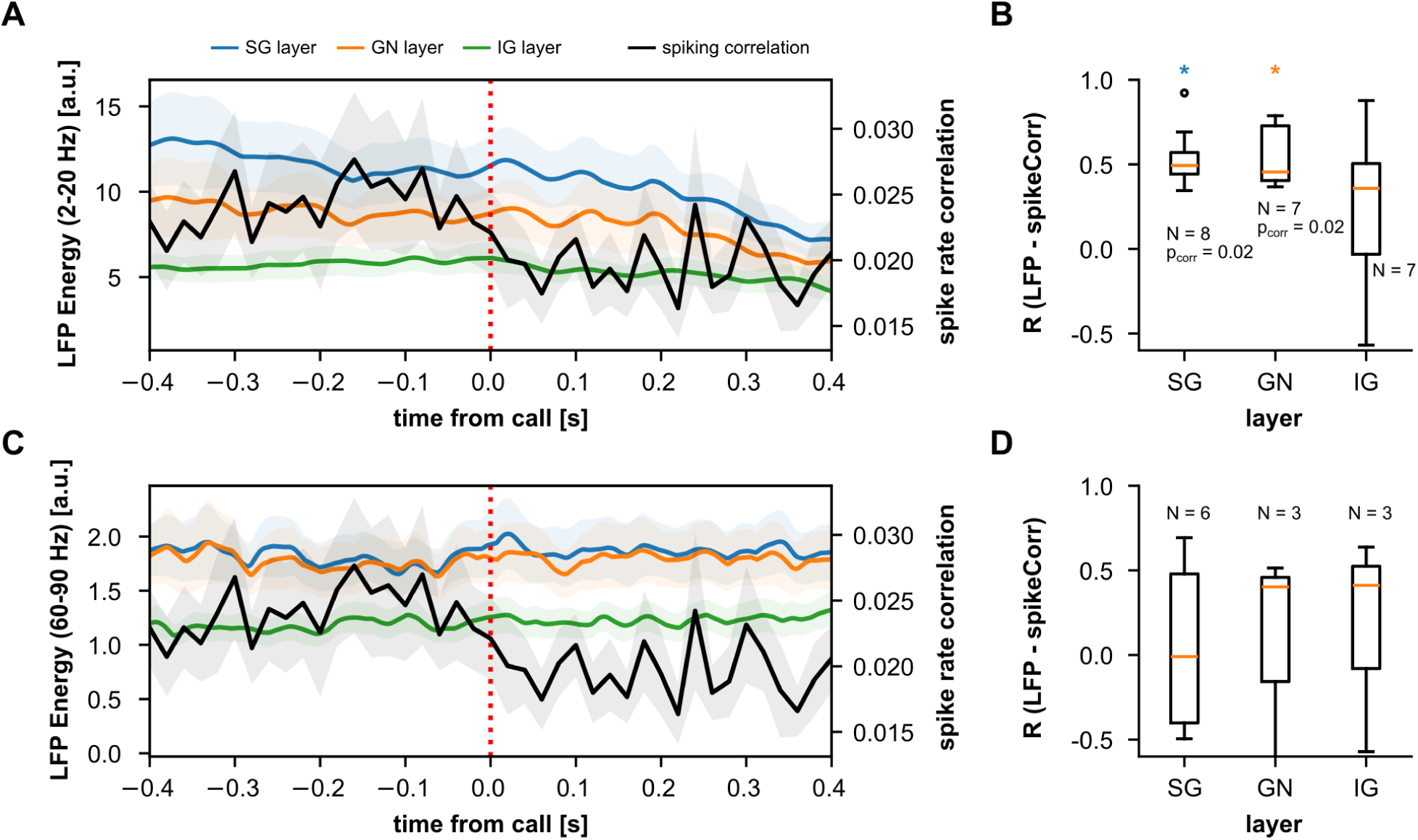
Spike-rate decorrelation and low-frequency LFP desynchronization are correlated over time. (**A**) Peri-vocal LFP energy (y-axis on the left) in low frequencies (12-20 Hz) for three cortical layer groups (SG, blue, N = 22 recordings; GN, orange, N = 22; IG, green, N = 18), and time-resolved spike-rate correlations (black, N = 22 recordings; y-axis on the right). Data shown as mean +- s.e.m. (**B**) Correlation (Spearman’s R) between spike-rate correlations and LFP energy in low-frequencies across layers. Only recordings with defined SG, GN or IG field potentials and which exhibited significant Spearman correlation (p < 0.05) were included in the values of each boxplot (N_SG_ = 8/22, 36%; N_GN_ = 7/22, 32%, N_IG_ = 7/18, 39%). Spearman’s R correlation values were significantly above 0 for SG and GN layers (p_corr_ < 0.05, indicated for each layer as a blue or orange star), but not layer IG (p_corr_ = 0.47). Note that correlation values were relatively high (median of 0.5). (**C**-**D**) Same as in **A**, **B** but LFP instantaneous energy values were obtained for high-frequency LFPs (gamma, 60-90 Hz). Note that the spike-rate correlation time-course is the same as that shown in **A**. Spearman’s R correlation values (**D**) were not significantly different than 0 for any layer.

**Fig S9.**
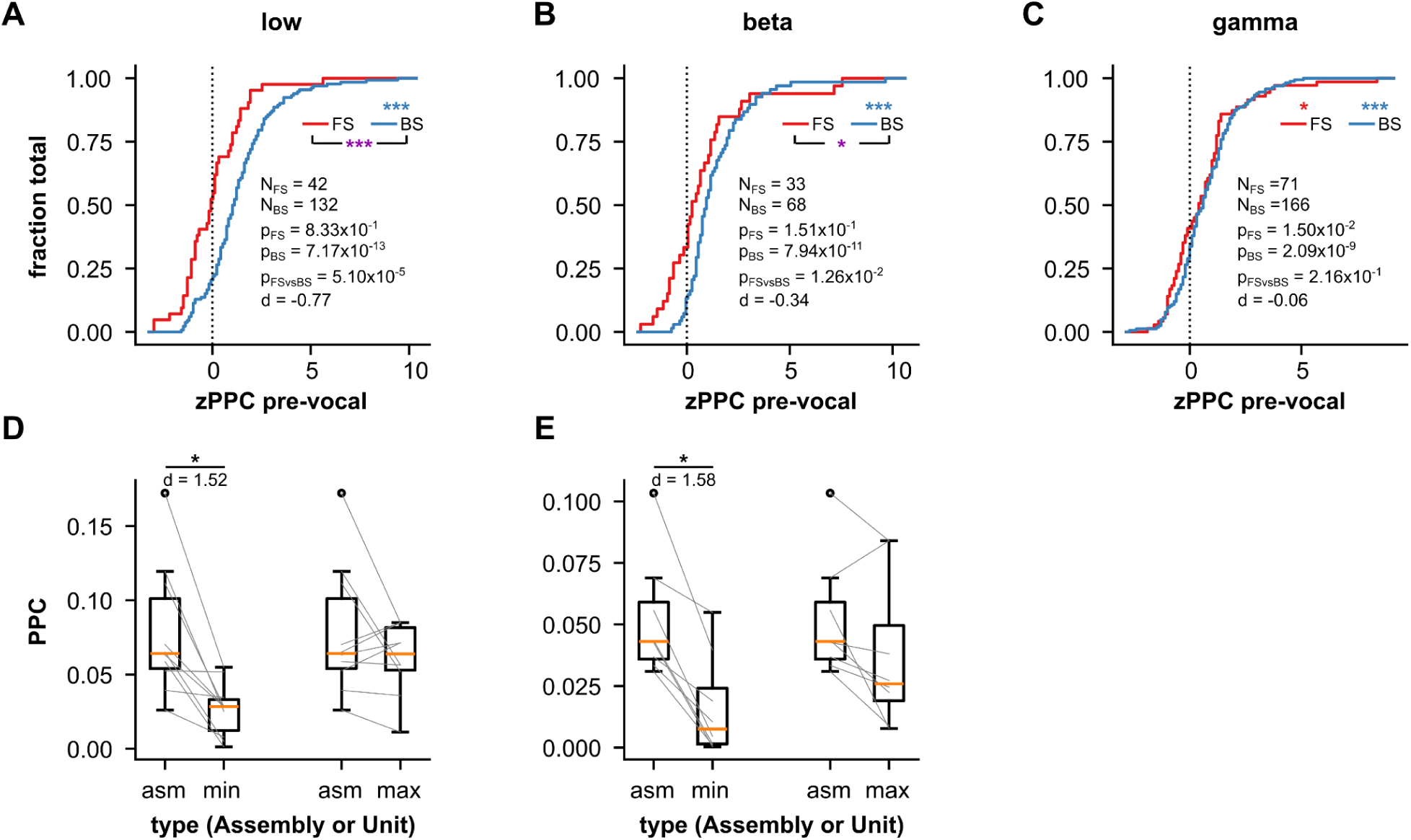
Pairwise-phase consistency (PPC) in AC during pre-vocal periods. (**A**) Cumulative distributions of z-normalised (relative to no-voc periods) PPC values computed between pre-vocal spiking and low-frequency (2-10 Hz) LFPs, shown for significantly (Rayleigh test, p < 0.05) synchronized FS (N_FS_ = 42; red) and BS (N_BS_ = 132, blue) units during the pre-vocal period. Z-normalised PPC was not significantly different than 0 across FS units (FDR-corrected Wilcoxon signed-rank test, p_FS_ = 0.83), but was significantly above 0 for BS units (p_BS_ = 7.17×10^-13^, blue ***). Normalised PPC values from FS units were significantly smaller than those from BS units (FDR-corrected Wilcoxon rank-sum test, p_FSvsBS_ = 5.10×10^-5^, purple ***) with moderate effect size (d = −0.77). (**B**) Same as **A**, but spike-LFP synchrony was calculated using beta-band (12-20 Hz) LFPs. (N_FS_ = 33, N_BS_ = 68; p_FS_ = 1.51×10^-1^, p_BS_ = 7.94×10^-11^; p_FSvsBS_ = 1.26×10^-2^, d = −0.34). (**C**) Same as **A**, but gamma-band LFPs were used for PPC calculation (N_FS_ = 71, N_BS_ = 166; p_FS_ = 1.50×10^-2^, p_BS_ = 2.09×10^-9^; p_FSvsBS_ = 2.16×10^-1^, d = −0.06). Red star indicates FS z-normalised FS values significantly different from 0. (**D**) Comparison between assembly PPC values (asm) vs. PPC values of constituent units with the smallest PPC (*min*) or the highest PPC (*max*), with spike-LFP synchrony computed for low-frequency LFPs during pre-vocal periods. Data is shown only for assemblies significantly synchronised to low-frequency LFPs (Rayleigh test, p < 0.05; N = 10). Assembly PPC values were significantly higher than those for its constituent *min* unit (FDR-corrected Wilcoxon signed-rank test, p_corr_ = 0.01) with large effect size (d = 1.52), but not for those of its constituent *max* unit (p_corr_ = 0.74). (**E**) Same as in D, but values related to beta-band LFPs are shown (N = 8). Assembly PPC values were significantly higher than those for its constituent *min* unit (p_corr_ = 0.02) with large effect size (d = 1.58), but not for those of its constituent *max* unit (p_corr_ = 0.54). In the figure: *, p_corr_ < 0.05; ***, p_corr_ < 0.001. Assembly PPC in the gamma-range were not shown because the number of significant assemblies was low (N = 1).

## Notes

### Competing Interest Statement

The authors have declared no competing interest.

## References

1. Hage, S. R. & Nieder, A. Dual Neural Network Model for the Evolution of Speech and Language. Trends Neurosci 39, 813–829 (2016).

2. Gavrilov, N., Hage, S. R. & Nieder, A. Functional Specialization of the Primate Frontal Lobe during Cognitive Control of Vocalizations. Cell Rep 21, 2393–2406 (2017).

3. Okobi, D. E., Jr., Banerjee, A., Matheson, A. M. M., Phelps, S. M. & Long, M. A. Motor cortical control of vocal interaction in neotropical singing mice. Science 363, 983–988 (2019).

4. Gavrilov, N. & Nieder, A. Distinct neural networks for the volitional control of vocal and manual actions in the monkey homologue of Broca’s area. eLife 10, e62797 (2021).

5. Babl, S. S., Kiai, A., García-Rosales, F. & Hechavarría, J. C. Neuronal activity underlying vocal production in bats. Ann N Y Acad Sci (2025) doi:10.1111/nyas.15410.

6. Suga, N., Schlegel, P., Shimozawa, T. & Simmons, J. Orientation sounds evoked from echolocating bats by electrical stimulation of the brain. J Acoust Soc Am 54, 793–797 (1973).

7. Fenzl, T. & Schuller, G. Periaqueductal gray and the region of the paralemniscal area have different functions in the control of vocalization in the neotropical bat, Phyllostomus discolor. Eur J Neurosci 16, 1974–1986 (2002).

8. Valentine, D. E., Sinha, S. R. & Moss, C. F. Orienting responses and vocalizations produced by microstimulation in the superior colliculus of the echolocating bat, Eptesicus fuscus. J Comp Physiol A Neuroethol Sens Neural Behav Physiol 188, 89–108 (2002).

9. Veerakumar, A., Head, J. P. & Krasnow, M. A. A brainstem circuit for phonation and volume control in mice. Nat Neurosci 26, 2122–2130 (2023).

10. Banerjee, A., Chen, F., Druckmann, S. & Long, M. A. Temporal scaling of motor cortical dynamics reveals hierarchical control of vocal production. Nat Neurosci 1–9 (2024) doi:10.1038/s41593-023-01556-5.

11. Smotherman, M. S. Sensory feedback control of mammalian vocalizations. Behavioural Brain Research 182, 315–326 (2007).

12. Eliades, S. J. & Wang, X. Neural substrates of vocalization feedback monitoring in primate auditory cortex. Nature 453, 1102–6 (2008).

13. Arriaga, G., Zhou, E. P. & Jarvis, E. D. Of Mice, Birds, and Men: The Mouse Ultrasonic Song System Has Some Features Similar to Humans and Song-Learning Birds. PLOS ONE 7, e46610 (2012).

14. Eliades, S. J. & Tsunada, J. Auditory cortical activity drives feedback-dependent vocal control in marmosets. Nat Commun 9, 2540 (2018).

15. Luo, J., Hage, S. R. & Moss, C. F. The Lombard Effect: From Acoustics to Neural Mechanisms. Trends Neurosci 41, 938–949 (2018).

16. Suga, N., Niwa, H., Taniguchi, I. & Margoliash, D. The personalized auditory cortex of the mustached bat: adaptation for echolocation. J Neurophysiol 58, 643–654 (1987).

17. Garcia-Rosales, F. et al. Echolocation-related reversal of information flow in a cortical vocalization network. Nat Commun 13, 3642 (2022).

18. Eliades, S. J. & Tsunada, J. Vocal error monitoring in the primate auditory cortex. J Neurosci e0090252025 (2025) doi:10.1523/JNEUROSCI.0090-25.2025.

19. Eliades, S. J. & Wang, X. Dynamics of auditory-vocal interaction in monkey auditory cortex. Cereb Cortex 15, 1510–1523 (2005).

20. Harmon, T. C., Madlon-Kay, S., Pearson, J. & Mooney, R. Vocalization modulates the mouse auditory cortex even in the absence of hearing. Cell Reports 43, 114611 (2024).

21. Schneider, D. M., Nelson, A. & Mooney, R. A synaptic and circuit basis for corollary discharge in the auditory cortex. Nature 513, 189–194 (2014).

22. Schneider, D. M. & Mooney, R. Motor-related signals in the auditory system for listening and learning. Curr Opin Neurobiol 33, 78–84 (2015).

23. Rummell, B. P., Klee, J. L. & Sigurdsson, T. Attenuation of Responses to Self-Generated Sounds in Auditory Cortical Neurons. J Neurosci 36, 12010–12026 (2016).

24. Li, S., Zhu, H. & Tian, X. Corollary Discharge Versus Efference Copy: Distinct Neural Signals in Speech Preparation Differentially Modulate Auditory Responses. Cereb Cortex 30, 5806–5820 (2020).

25. Eliades, S. J. & Wang, X. Sensory-motor interaction in the primate auditory cortex during self-initiated vocalizations. J Neurophysiol 89, 2194–2207 (2003).

26. Linden, J. F. & Schreiner, C. E. Columnar transformations in auditory cortex? A comparison to visual and somatosensory cortices. Cereb Cortex 13, 83–9 (2003).

27. Park, Y. & Geffen, M. N. A circuit model of auditory cortex. PLOS Computational Biology 16, e1008016 (2020).

28. Gungor Aydin, A., Lemenze, A. & Bieszczad, K. M. Functional diversities within neurons and astrocytes in the adult rat auditory cortex revealed by single-nucleus RNA sequencing. Sci Rep 14, 25314 (2024).

29. Studer, F. & Barkat, T. R. Inhibition in the auditory cortex. Neuroscience & Biobehavioral Reviews 132, 61–75 (2022).

30. Gillet, S. N., Kato, H. K., Justen, M. A., Lai, M. & Isaacson, J. S. Fear Learning Regulates Cortical Sensory Representations by Suppressing Habituation. Front Neural Circuits 11, 112 (2017).

31. Kuchibhotla, K. V. et al. Parallel processing by cortical inhibition enables context-dependent behavior. Nat Neurosci 20, 62–71 (2017).

32. Liang, F. et al. Sparse Representation in Awake Auditory Cortex: Cell-type Dependence, Synaptic Mechanisms, Developmental Emergence, and Modulation. Cereb Cortex 29, 3796–3812 (2019).

33. Fernandez, A. A., Fasel, N., Knörnschild, M. & Richner, H. When bats are boxing: aggressive behaviour and communication in male Seba’s short-tailed fruit bat. Animal Behaviour 98, 149–156 (2014).

34. Knörnschild, M., Feifel, M. & Kalko, E. K. V. Male courtship displays and vocal communication in the polygynous bat Carollia perspicillata. (2014) doi:10.1163/1568539X-00003171.

35. Hechavarria, J. C., Beetz, M. J., Macias, S. & Kossl, M. Distress vocalization sequences broadcasted by bats carry redundant information. J Comp Physiol A Neuroethol Sens Neural Behav Physiol 202, 503–15 (2016).

36. Williams, C. F. Social Organization of the Bat, Carollia perspicillata (Chiroptera: Phyllostomidae). Ethology 71, 265–282 (1986).

37. Porter, F. L. Social Behavior in the Leaf-Nosed Bat, Carollia perspicillata. Zeitschrift für Tierpsychologie 50, 1–8 (1979).

38. Hechavarria, J. C., Beetz, M. J., Macias, S. & Kossl, M. Vocal sequences suppress spiking in the bat auditory cortex while evoking concomitant steady-state local field potentials. Sci Rep 6, 39226 (2016).

39. Buzsáki, G. Neural syntax: cell assemblies, synapsembles and readers. Neuron 68, 362–385 (2010).

40. Russo, E. & Durstewitz, D. Cell assemblies at multiple time scales with arbitrary lag constellations. Elife 6, e19428 (2017).

41. Sotomayor-Gómez, B., Battaglia, F. P. & Vinck, M. SpikeShip: A method for fast, unsupervised discovery of high-dimensional neural spiking patterns. PLoS Comput Biol 19, e1011335 (2023).

42. Sotomayor-Gómez, B., Battaglia, F. P. & Vinck, M. Firing rates in visual cortex show representational drift, while temporal spike sequences remain stable. Cell Reports 44, (2025).

43. Buzsaki, G., Anastassiou, C. A. & Koch, C. The origin of extracellular fields and currents--EEG, ECoG, LFP and spikes. Nat Rev Neurosci 13, 407–20 (2012).

44. Lakatos, P. et al. The Spectrotemporal Filter Mechanism of Auditory Selective Attention. Neuron 77, 750–761 (2013).

45. Tavares, L. C. S. & Tort, A. B. L. Hippocampal-prefrontal interactions during spatial decision-making. Hippocampus 32, 38–54 (2022).

46. Uran, C. et al. Predictive coding of natural images by V1 firing rates and rhythmic synchronization. Neuron 110, 2886–2887 (2022).

47. Jacobs, E. A. K., Steinmetz, N. A., Peters, A. J., Carandini, M. & Harris, K. D. Cortical State Fluctuations during Sensory Decision Making. Current Biology 30, 4944–4955.e7 (2020).

48. Vinck, M., van Wingerden, M., Womelsdorf, T., Fries, P. & Pennartz, C. M. The pairwise phase consistency: a bias-free measure of rhythmic neuronal synchronization. Neuroimage 51, 112–22 (2010).

49. García-Rosales, F., Schaworonkow, N. & Hechavarria, J. C. Oscillatory Waveform Shape and Temporal Spike Correlations Differ across Bat Frontal and Auditory Cortex. J Neurosci 44, e1236232023 (2024).

50. Oberto, V. J. et al. Distributed cell assemblies spanning prefrontal cortex and striatum. Curr Biol 32, 1–13.e6 (2022).

51. Müller-Preuss, P. & Ploog, D. Inhibition of auditory cortical neurons during phonation. Brain Res 215, 61–76 (1981).

52. Tsunada, J. & Eliades, S. J. Dissociation of Unit Activity and Gamma Oscillations during Vocalization in Primate Auditory Cortex. J. Neurosci. 40, 4158–4171 (2020).

53. Eliades, S. J. & Wang, X. Contributions of sensory tuning to auditory-vocal interactions in marmoset auditory cortex. Hearing Research 348, 98–111 (2017).

54. Flinker, A. et al. Single-Trial Speech Suppression of Auditory Cortex Activity in Humans. J Neurosci 30, 16643–16650 (2010).

55. Bigelow, J., Morrill, R. J., Dekloe, J. & Hasenstaub, A. R. Movement and VIP Interneuron Activation Differentially Modulate Encoding in Mouse Auditory Cortex. eNeuro 6, ENEURO.0164-19.2019 (2019).

56. Ozker, M. et al. Speech-induced suppression and vocal feedback sensitivity in human cortex. Elife 13, RP94198 (2024).

57. Nelson, A. et al. A Circuit for Motor Cortical Modulation of Auditory Cortical Activity. J. Neurosci. 33, 14342–14353 (2013).

58. Sakata, S. & Harris, K. D. Laminar-dependent effects of cortical state on auditory cortical spontaneous activity. Front. Neural Circuits 6, (2012).

59. McGinley, M. J., David, S. V. & McCormick, D. A. Cortical Membrane Potential Signature of Optimal States for Sensory Signal Detection. Neuron 87, 179–192 (2015).

60. Schwartz, Z. P., Buran, B. N. & David, S. V. Pupil-associated states modulate excitability but not stimulus selectivity in primary auditory cortex. J Neurophysiol 123, 191–208 (2020).

61. Otazu, G. H., Tai, L.-H., Yang, Y. & Zador, A. M. Engaging in an auditory task suppresses responses in auditory cortex. Nat Neurosci 12, 646–654 (2009).

62. Niell, C. M. & Stryker, M. P. Modulation of Visual Responses by Behavioral State in Mouse Visual Cortex. Neuron 65, 472–479 (2010).

63. Vinck, M., Batista-Brito, R., Knoblich, U. & Cardin, J. A. Arousal and Locomotion Make Distinct Contributions to Cortical Activity Patterns and Visual Encoding. Neuron 86, 740–754 (2015).

64. Harris, K. D. & Thiele, A. Cortical state and attention. Nat Rev Neurosci 12, 509–523 (2011).

65. Nelson, A. & Mooney, R. The Basal Forebrain and Motor Cortex Provide Convergent yet Distinct Movement-Related Inputs to the Auditory Cortex. Neuron 90, 635–648 (2016).

66. Borjon, J. I., Takahashi, D. Y., Cervantes, D. C. & Ghazanfar, A. A. Arousal dynamics drive vocal production in marmoset monkeys. J Neurophysiol 116, 753–764 (2016).

67. Jürgens, U. Vocalization as an emotional indicator. A neuroethological study in the squirrel monkey. Behaviour 69, 88–117 (1979).

68. Liao, D. A., Zhang, Y. S., Cai, L. X. & Ghazanfar, A. A. Internal states and extrinsic factors both determine monkey vocal production. Proceedings of the National Academy of Sciences 115, 3978–3983 (2018).

69. Nieder, A. & Mooney, R. The neurobiology of innate, volitional and learned vocalizations in mammals and birds. Philosophical Transactions of the Royal Society B: Biological Sciences 375, 20190054 (2019).

70. Kobler, J. B., Isbey, S. F. & Casseday, J. H. Auditory pathways to the frontal cortex of the mustache bat, Pteronotus parnellii. Science 236, 824–6 (1987).

71. Casseday, J. H., Kobler, J. B., Isbey, S. F. & Covey, E. Central acoustic tract in an echolocating bat: an extralemniscal auditory pathway to the thalamus. J Comp Neurol 287, 247–59 (1989).

72. Nevue, A. A., Mello, C. V. & Portfors, C. V. Bats possess the anatomical substrate for a laryngeal motor cortex*. bioRxiv* 2023.06.26.546619 (2023) doi:10.1101/2023.06.26.546619.

73. Tsunada, J., Wang, X. & Eliades, S. J. Multiple processes of vocal sensory-motor interaction in primate auditory cortex. Nat Commun 15, 3093 (2024).

74. Letzkus, J. J. et al. A disinhibitory microcircuit for associative fear learning in the auditory cortex. Nature 480, 331–335 (2011).

75. Harris, K. D., Csicsvari, J., Hirase, H., Dragoi, G. & Buzsáki, G. Organization of cell assemblies in the hippocampus. Nature 424, 552–556 (2003).

76. Miehl, C., Onasch, S., Festa, D. & Gjorgjieva, J. Formation and computational implications of assemblies in neural circuits. The Journal of Physiology 601, 3071–3090 (2023).

77. Bocchio, M. et al. Functional networks of inhibitory neurons orchestrate synchrony in the hippocampus. PLOS Biology 22, e3002837 (2024).

78. Garcia-Rosales, F., Beetz, M. J., Cabral-Calderin, Y., Kossl, M. & Hechavarria, J. C. Neuronal coding of multiscale temporal features in communication sequences within the bat auditory cortex. Commun Biol 1, 200 (2018).

79. Kayser, C., Montemurro, M. A., Logothetis, N. K. & Panzeri, S. Spike-Phase Coding Boosts and Stabilizes Information Carried by Spatial and Temporal Spike Patterns. Neuron 61, 597–608 (2009).

80. Luczak, A., McNaughton, B. L. & Harris, K. D. Packet-based communication in the cortex. Nat Rev Neurosci 16, 745–755 (2015).

81. Waschke, L., Tune, S. & Obleser, J. Local cortical desynchronization and pupil-linked arousal differentially shape brain states for optimal sensory performance. eLife 8, e51501 (2019).

82. Marguet, S. L. & Harris, K. D. State-dependent representation of amplitude-modulated noise stimuli in rat auditory cortex. J Neurosci 31, 6414–6420 (2011).

83. Pachitariu, M., Lyamzin, D. R., Sahani, M. & Lesica, N. A. State-Dependent Population Coding in Primary Auditory Cortex. J. Neurosci. 35, 2058–2073 (2015).

84. Treviño, M. Inhibition Controls Asynchronous States of Neuronal Networks. Front. Synaptic Neurosci. 8, (2016).

85. Giraud, A.-L. & Poeppel, D. Cortical oscillations and speech processing: emerging computational principles and operations. Nat Neurosci 15, 511–517 (2012).

86. Garcia-Rosales, F. et al. Low-Frequency Spike-Field Coherence Is a Fingerprint of Periodicity Coding in the Auditory Cortex. iScience 9, 47–62 (2018).

87. Gregoriou, G. G., Gotts, S. J., Zhou, H. & Desimone, R. High-frequency, long-range coupling between prefrontal and visual cortex during attention. Science 324, 1207–1210 (2009).

88. Vinck, M., Womelsdorf, T., Buffalo, E. A., Desimone, R. & Fries, P. Attentional modulation of cell-class specific gamma-band synchronization in awake monkey area V4. Neuron 80, 10.1016/j.neuron.2013.08.019 (2013).

89. Sellers, K. K. et al. Oscillatory dynamics in the frontoparietal attention network during sustained attention in the ferret. Cell Rep 16, 2864–2874 (2016).

90. Schneider, M. et al. A mechanism for inter-areal coherence through communication based on connectivity and oscillatory power. Neuron 109, 4050–4067.e12 (2021).

91. Beetz, M. J., Hechavarría, J. C. & Kössl, M. Temporal tuning in the bat auditory cortex is sharper when studied with natural echolocation sequences. Sci Rep 6, 29102 (2016).

92. Garcia-Rosales, F., Lopez-Jury, L., Gonzalez-Palomares, E., Cabral-Calderin, Y. & Hechavarria, J. C. Fronto-Temporal Coupling Dynamics During Spontaneous Activity and Auditory Processing in the Bat Carollia perspicillata. Front Syst Neurosci 14, 14 (2020).

93. Eiermann, A. & Esser, K. H. Auditory responses from the frontal cortex in the short-tailed fruit bat Carollia perspicillata. Neuroreport 11, 421–5 (2000).

94. Garcia-Rosales, F. et al. Laminar specificity of oscillatory coherence in the auditory cortex. Brain Struct Funct 224, 2907–2924 (2019).

95. Buccino, A. P. et al. SpikeInterface, a unified framework for spike sorting. Elife 9, e61834 (2020).

96. Pachitariu, M., Sridhar, S., Pennington, J. & Stringer, C. Spike sorting with Kilosort4. Nat Methods 21, 914–921 (2024).

97. Ardid, S. et al. Mapping of Functionally Characterized Cell Classes onto Canonical Circuit Operations in Primate Prefrontal Cortex. J. Neurosci. 35, 2975–2991 (2015).

98. Montijn, J. S. et al. A parameter-free statistical test for neuronal responsiveness. Elife 10, (2021).

99. Stringer, C. et al. Rastermap: a discovery method for neural population recordings. Nat Neurosci 28, 201–212 (2025).

100. Maris, E. & Oostenveld, R. Nonparametric statistical testing of EEG- and MEG-data. J Neurosci Methods 164, 177–190 (2007).

101. Denker, M., Yegenoglu, A. & Grün, S. Collaborative HPC-enabled workflows on the HBP Collaboratory using the Elephant framework. in Neuroinformatics 2018 P19 (2018). doi:10.12751/incf.ni2018.0019.

102. Cole, S., Donoghue, T., Gao, R. & Voytek, B. %J J. of O. S. S. NeuroDSP: a package for neural digital signal processing. 4, 1272 (2019).

103. Astropy Collaboration, et al. Astropy: A community Python package for astronomy. åp 558, A33 (2013).

104. Donoghue, T. et al. Parameterizing neural power spectra into periodic and aperiodic components. Nat Neurosci 23, 1655–1665 (2020).

105. Benjamini, Y. & Hochberg, Y. Controlling the False Discovery Rate: A Practical and Powerful Approach to Multiple Testing. Journal of the Royal Statistical Society: Series B (Methodological) 57, 289–300 (1995).

106. Cohen, J. Statistical Power Analysis for the Behavioral Sciences. (Routledge, 2013).

