## Supplementary Figures S1-S9 for "Vocal production differentially affects fast- and broad-spiking neurons in auditory cortex"

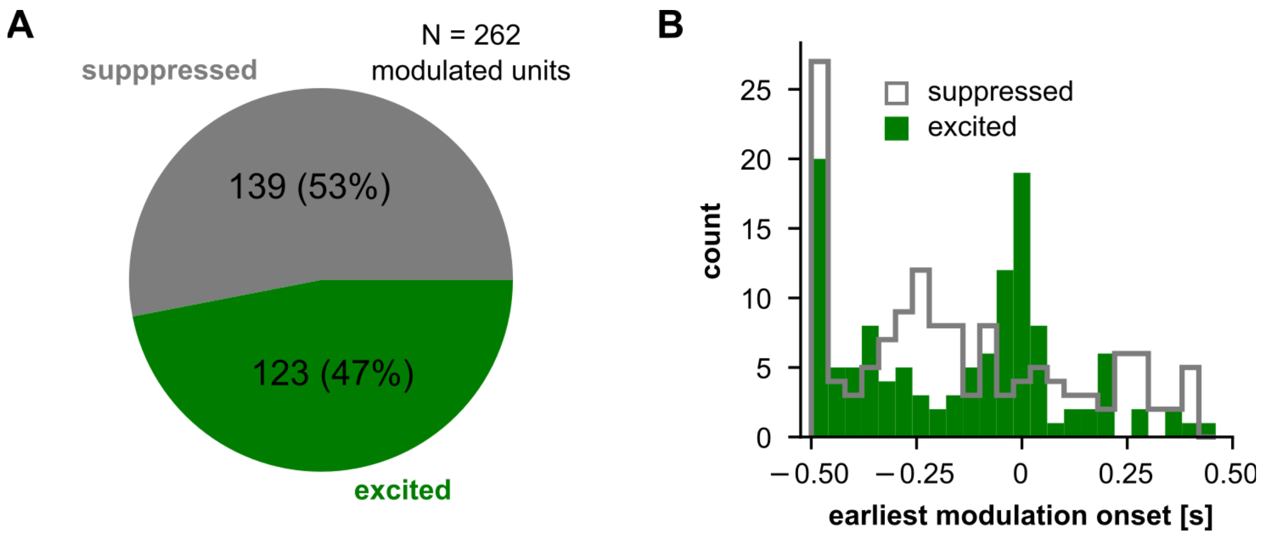


**Fig. S1. Distribution of vocalisation-related modulations in AC.**

(**A**) Proportion of suppressed and excited units in the vocalisation-related populations. (**B**) Distribution of earliest modulation onset (the onset of the first significantly modulated cluster across units) for suppression and excitation.

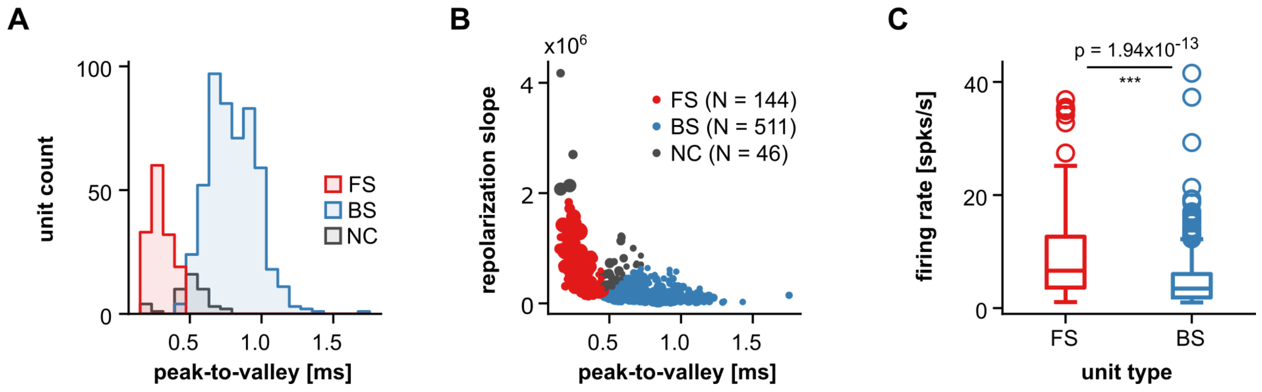


**Fig. S2. Spiking properties of FS and BS units.**

(**A**) Distribution of peak-to-valley durations across neuron types. (**B**) Peak-to-valley and repolarisation slopes for all units (N = 701). Each circle represents a unit, with size depending on its firing rate. A k-mean clustering using these two variables, together with a 460 µs threshold for peak-to-valley duration (as in **A**) were used for unit classification (see Methods). (**C**) The firing rates of FS units were significantly higher than those of their BS counterparts (Wilcoxon rank-sum test, $p = 1.94 \times 10^{-13}$).

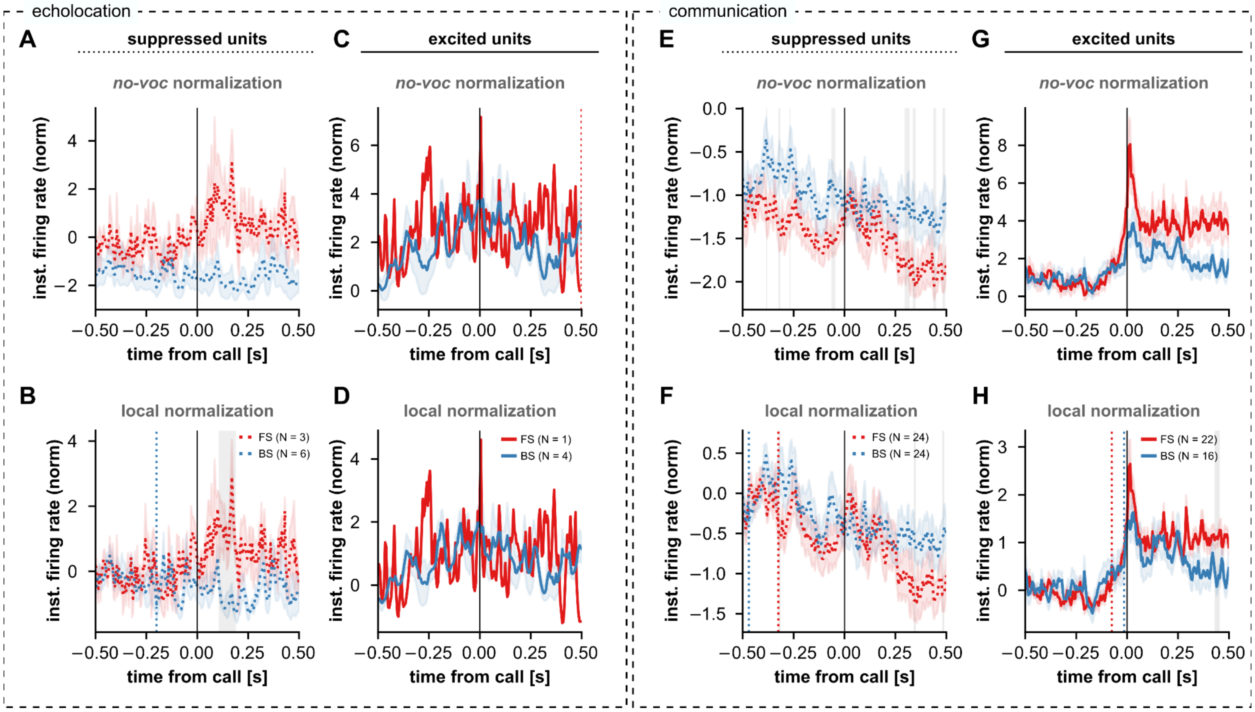


**Fig. S3. Call-type specific, peri-vocal firing patterns of FS- and BS units.**

(**A-D**) Follows **Fig. 3A-D**, but showing peri-vocal firing patterns of significantly modulated FS and BS units when animals emitted echolocation calls. (**E-H**) Same as **A-D**, but peri-vocal spiking patterns were considered only when animals vocalised communication calls. The sample sizes were not sufficient to draw conclusions from analysing peri-vocal activity in a call-specific manner, even when relaxing the minimum number of trials necessary for a unit to be considered (here: at least 10 vocalisations; in the main analysis: at least 15). For example, in the echolocation case only 1 excited FS cell was found (panels **C, D**). The reason is the sparseness of echolocation calling in our sessions. Indeed, most echolocation utterances were found in a small number of recordings with relatively low neuronal yield. In other recordings, echolocation utterances contributed to the trial pool, but were not enough to be analysed on their own on a neuron-per-neuron basis. Therefore, and in order to maximise the number of vocalisation trials and the number of vocalisation-modulated units considered, echolocation and communication calls were both considered as “vocalisations” in the main text.

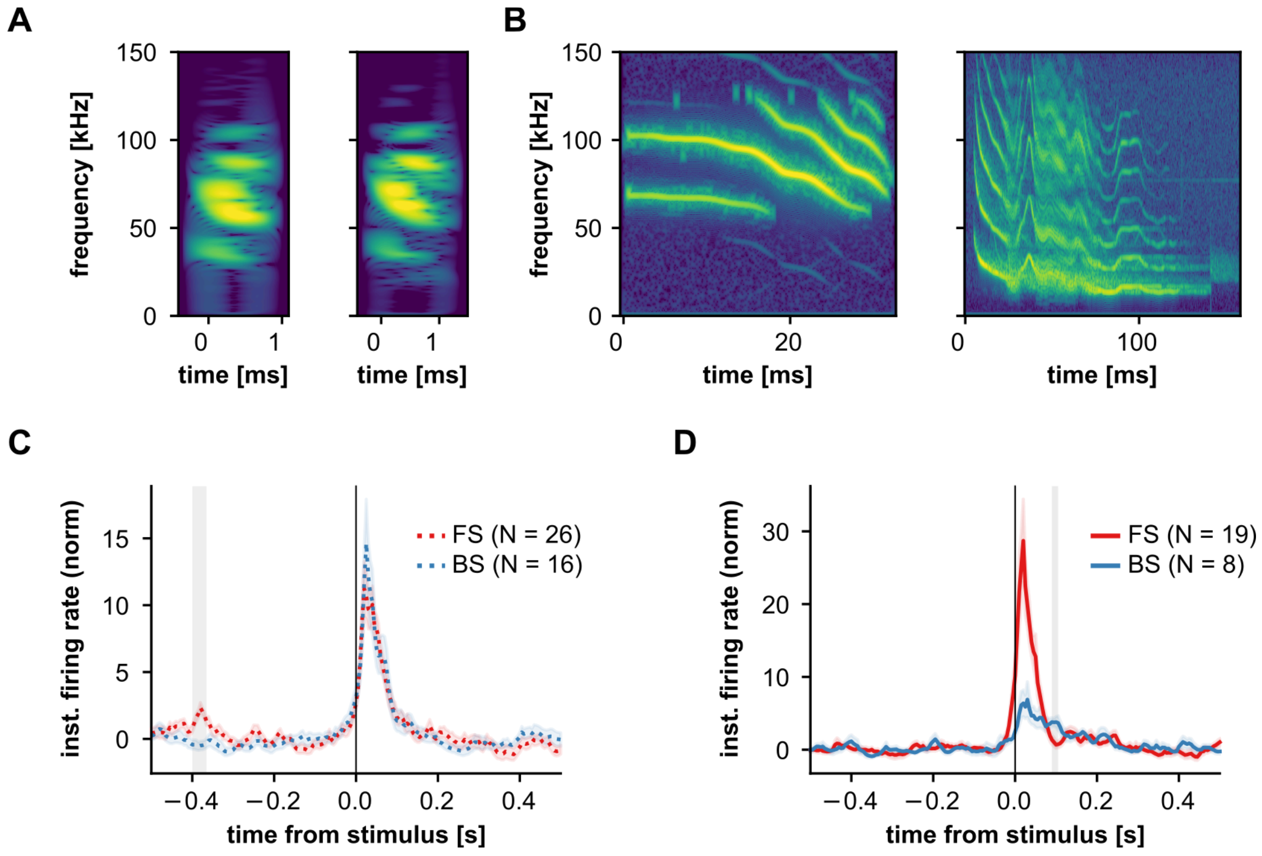


**Fig. S4. Spiking activity of vocalisation-modulated units in response to acoustic stimulation with natural calls.**

(**A, B**) Natural stimuli used for acoustic stimulation: two echolocation (**A**) and two communication (**B**) calls. (**C**) IFRs of significantly stimulus-responsive (cluster-based statistics) FS (red; N = 26) and BS (blue) units, normalised to a no-voc baseline. These units were in addition significantly suppressed during vocal production. Firing rates were not significantly different throughout the stimulation period (cluster-based statistics), except for a significant time segment shown as a gray area at about -400 ms. Without stimulation at those time-points, the interpretation for this significant cluster is unclear. (**D**) Same as **C**, but considering vocalisation-related excited FS and BS units. Apparent differences between FS and BS units immediately after vocalisation onset were not significant (cluster-based statistics), possibly due to firing rate variability and relatively small sample sizes (FS, N = 19; BS, N = 8). In both **C** and **D**, significant stimulus responsiveness was determined using cluster-based statistics similar to those used to determine significant vocalisation-related modulations. Differences for the IFRs across cell types were statistically assessed using the same paradigm as that of **Fig. 3**.

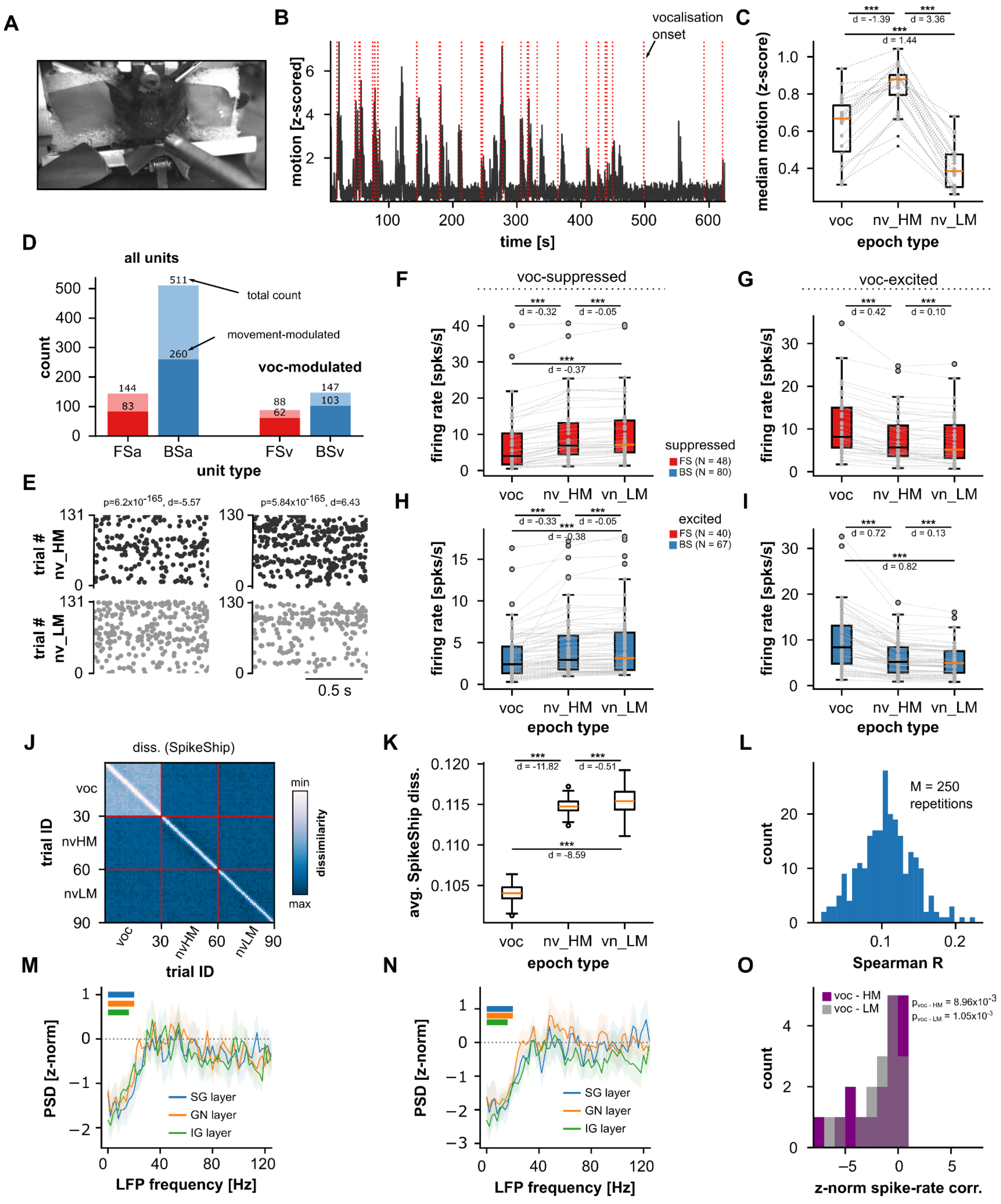


**Fig S5. Movement alone does not explain vocalisation-related neural dynamics.**

(**A**) A still-frame corresponding to one session's video recording. (**B**) Motion quantification for a 10-minute sub-session in a representative recording. Vocalisation onsets are marked with red dashed lines. Vocalisation is paired with movement, but movement also occurs at times when no calls are uttered. (**C**) Comparison of motion in three epoch types, across all recording sessions (N = 22): vocalisation (voc), no-voc high-motion (nv_HM; i.e. no-voc epochs with the highest 30th percentile of quantified motion), and no-voc low-motion (nv_LM; no-voc epochs with the lowest 30th percentile of quantified motion). Movement during vocalisation was significantly higher (with large effect size) than during nv_LM periods (Wilcoxon signed-rank test, $p = 9.54x10^{-7}$, d = 1.44). Note, however, that movement during nv_HM epochs was the highest of all epoch types (also than vocalisation), with large effect sizes ($p \leq 7.1x10^{-7}$, $|d| \geq 1.39$). (**D**) Number of units in the AC, modulated by the animal's movement, according to cell type (FS, red; BS, blue). *Left*: considering all of the units; *right*: considering only units that were significantly modulated by vocalisation. Total and relative numbers are indicated in the figure. 58% of all FS units (and 70% of vocalisation-modulated FS units) were modulated by movement. 50% of all BS units (and 70% of vocalisation-modulated BS units) were modulated by movement. A unit was considered modulated by movement if the firing rate in nv_HM epochs was different than in nv_LM ones (Wilcoxon rank-sum test, $p < 0.05$), with at least moderate effect sizes ($|d| >= 0.5$). (**E**) *Left*: Raster plots for an example movement-modulated unit (top, nv_HM, dark spikes; bottom: nv_LM, gray spikes). One repetition of nv_HM and nv_LM bootstrap iteration is shown (see Method). Note that this unit's firing rate decreased in nv_HM epochs (indicated by negative d, shown on top). *Right*: A similar representation, but for an exemplary unit whose firing rate increased during nv_HM trials relative to nv_LM ones. (**F, H**) Firing rates for vocalisation-suppressed cells during voc, nv_HM, and nv_LM epochs, shown for FS (**F**) and BS (**H**) units. (**G, I**) Same as in **F,H**, but vocalisation-excited units are shown. These units are the same used in **Fig. 3** of the main text. Vocalisation-related modulations were always significantly stronger (either for suppression or excitation) than in either high- or low-movement no-voc epochs (Wilcoxon signed-rank test, $p \leq 2.45x10^{-9}$, $|d| \geq 0.32$). Note that this occurs even when movement in nv_HM was stronger than in vocalisation epochs. Thus, firing rate modulations observed in our dataset are not wholly accounted for by the animal's motion. (**J**) Median SpikeShip dissimilarity matrix (see **Fig. 4**) considering epochs of vocalisation, nv_HM, and nv_LM. (**K**) Spike temporal patterns were significantly more stable during vocalisation epochs than during either high- or low- movement epochs, with large effect sizes (Wilcoxon rank-sum test, $p \leq 3.33x10^{-83}$, $d < -8.6$). (**L**) Distribution of the correlation between SpikeShip and and firing rate dissimilarity matrices, across repetitions (M = 250). Relatively low correlation values indicate independence between the temporal patterns and the neuronal firing rates. Altogether, panels **J-L** indicate that the neuronal temporal structure observed during vocalisation (**Fig. 4**) cannot be fully accounted for by movement. (**M**) Z-normalised pre-vocal LFP power relative to hv_HM periods across cortical layers. Horizontal lines at the top with colours matching layer identifying colours, indicate frequency ranges for which z-normalised power values across recordings are significantly different than 0 (FDR-corrected Wilcoxon signed-rank tests across frequencies, $p_{corr} < 0.05$). (**N**) Same as in **M**, but pre-vocal LFP power was normalised to nv_LM epochs. (**O**) Distribution of z-normalised spike firing rate correlations during peri-vocal epochs, relative to nv_HM (purple) or nv_LM (gray). Normalised spiking correlations were significantly below zero in both cases (Wilcoxon signed-rank test, $p \leq 8.96x10^{-3}$, N = 18 recordings). Panels **M-O** indicate that the neural decorrelation, observed in the LFP and the spiking activity (**Fig. 5**), cannot be fully explained by the animal's movement.

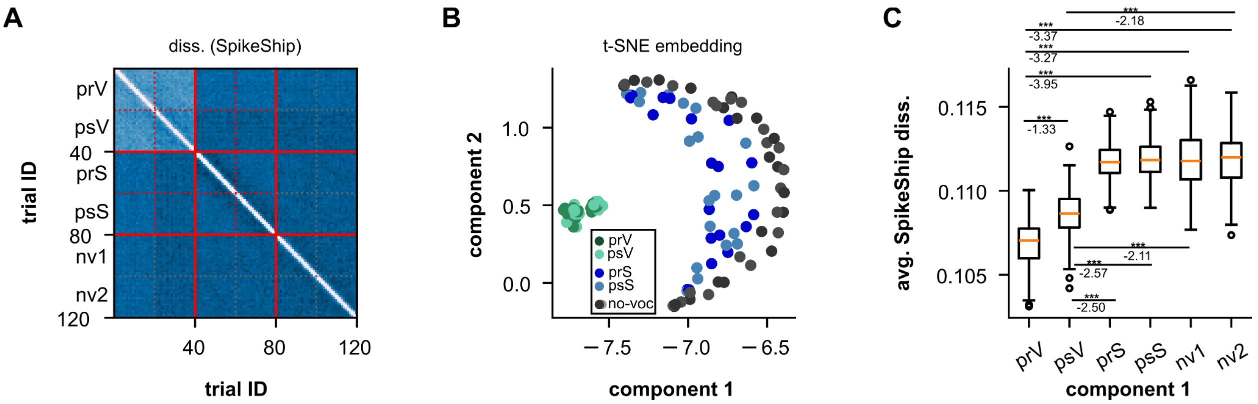


**Fig. S6. SpikeShip pattern analysis including epochs of acoustic stimulation with natural stimuli.**

(**A**) Average SpikeShip dissimilarity matrix across repetitions (M = 250), for epochs that include pre-vocal, post-vocal, pre-stimulus, post-stimulus, and no-voc trial types. The SpikeShip analysis is similar to the one of **Fig. 4** in the main text. However, 20 epochs (instead of 30) of each type were selected, and units were only included if they had sufficient vocalisation and passive listening trials (i.e. at least 20). Because of these requisites, only a subset of 282 units were used (in the main analysis, 417 were used). (**B**) 2-dimensional t-SNE embedding obtained from the dissimilarity matrix in **A**. (**C**) Average SpikeShip dissimilarity across trial types. Values for pre-vocal and post-vocal epochs were significantly smaller than for pre-/post-stimulus and no-voc trials with large effect sizes (FDR-Corrected Wilcoxon rank-sum tests, $p_{corr} < 9.80 \times 10^{-63}$; $d < -2.11$). SpikeShip dissimilarities were also different for pre-vocal and post-vocal trials ($p_{corr} < 3.42 \times 10^{-37}$; $d < -1.33$). Altogether, these results support that spike temporal patterns formed during vocal production are unique to this behavioural state.

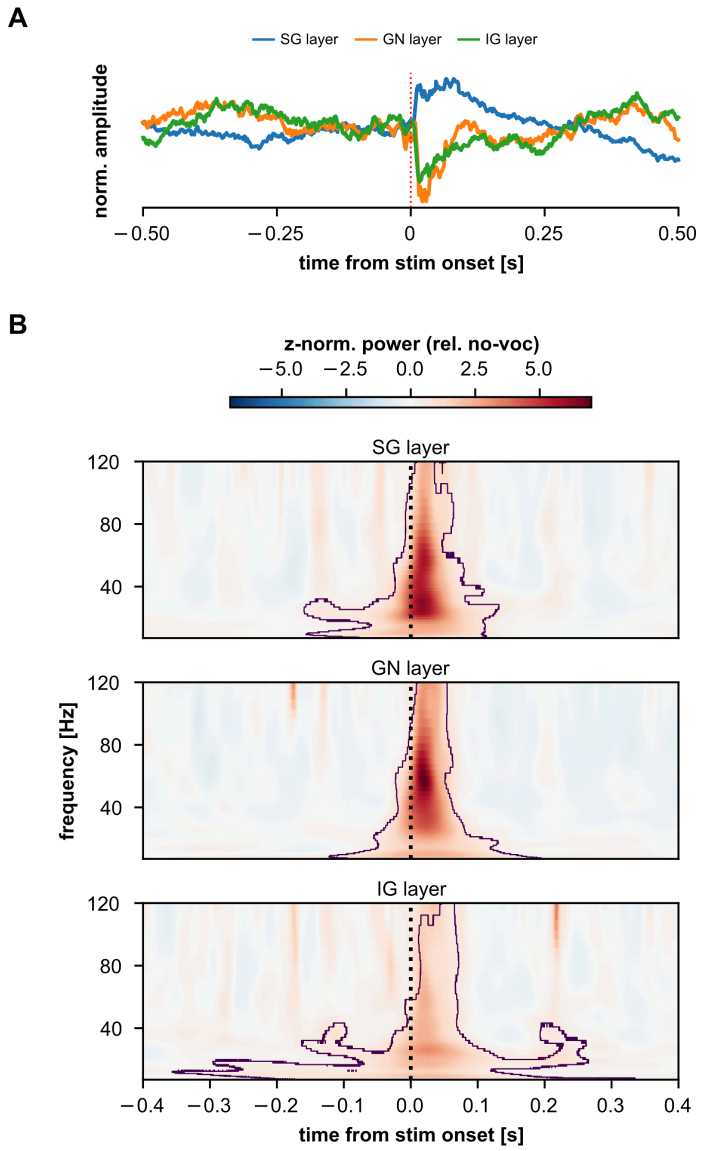


**Fig. S7. LFP response to acoustic stimulation.**

(**A**) Stimulus-evoked LFPs from a representative recording session (SG: supragranular LFPs, GN: granular LFPs, IG: infragranular LFPs). (**B**) Stimulus-evoked time-frequency resolved LFP spectra across cortical layers, averaged across recording sessions. The treatment to these spectra is the same as that shown in **Fig. 5D**: they were z-normalised to a no-voc baseline. Significant deviations from 0 across recordings were interpreted as a consistent trend in our dataset. Contour lines in the figure delineate significant clusters (see Methods). A clear evoked-related pattern indicates typical primary-like auditory responses to the natural stimuli. No evidence for low-frequency LFP desynchronization is visible during passive listening (cf. **Fig. 5**).

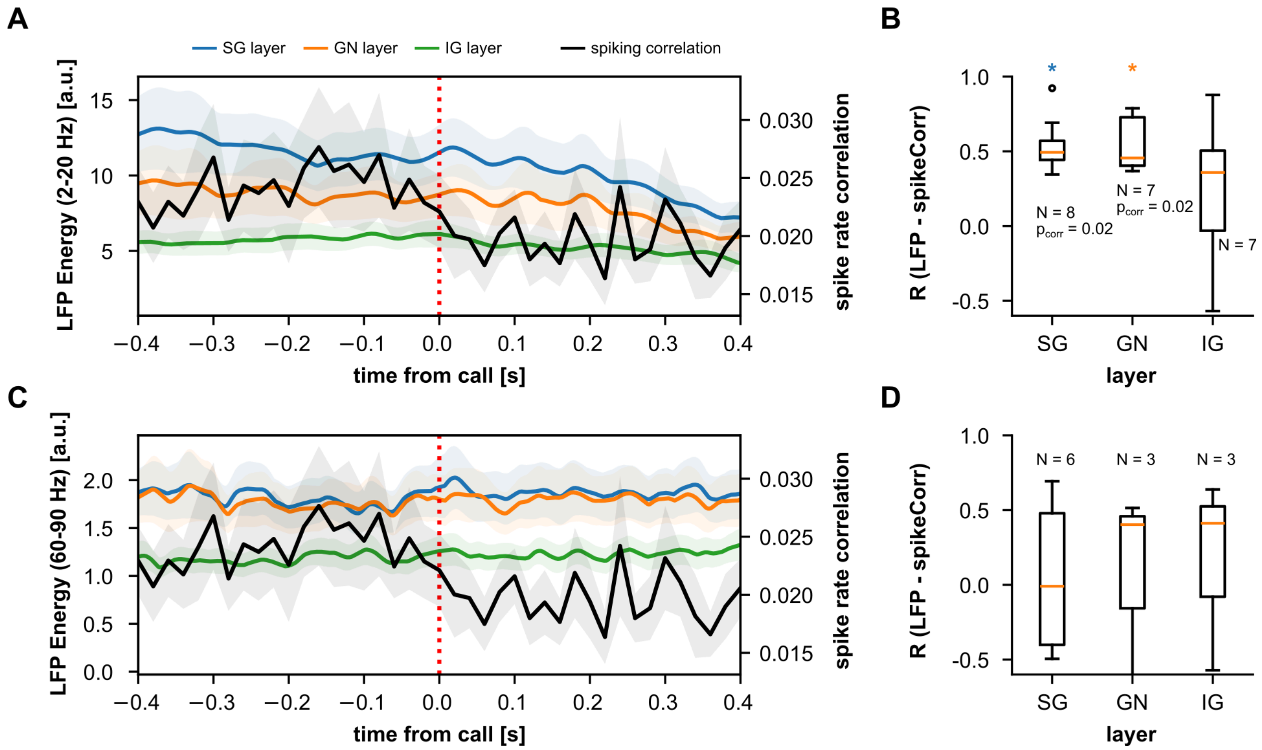


**Fig S8. Spike-rate decorrelation and low-frequency LFP desynchronization are correlated over time.**

(**A**) Peri-vocal LFP energy (y-axis on the left) in low frequencies (12-20 Hz) for three cortical layer groups (SG, blue, N = 22 recordings; GN, orange, N = 22; IG, green, N = 18), and time-resolved spike-rate correlations (black, N = 22 recordings; y-axis on the right). Data shown as mean +- s.e.m. (**B**) Correlation (Spearman's R) between spike-rate correlations and LFP energy in low-frequencies across layers. Only recordings with defined SG, GN or IG field potentials and which exhibited significant Spearman correlation ($p < 0.05$) were included in the values of each boxplot ($N_{SG}$ = 8/22, 36%; $N_{GN}$ = 7/22, 32%, $N_{IG}$ = 7/18, 39%). Spearman's R correlation values were significantly above 0 for SG and GN layers ($p_{corr} < 0.05$, indicated for each layer as a blue or orange star), but not layer IG ($p_{corr} = 0.47$). Note that correlation values were relatively high (median of 0.5). (**C**-**D**) Same as in **A**, **B** but LFP instantaneous energy values were obtained for high-frequency LFPs (gamma, 60-90 Hz). Note that the spike-rate correlation time-course is the same as that shown in **A**. Spearman's R correlation values (**D**) were not significantly different than 0 for any layer.

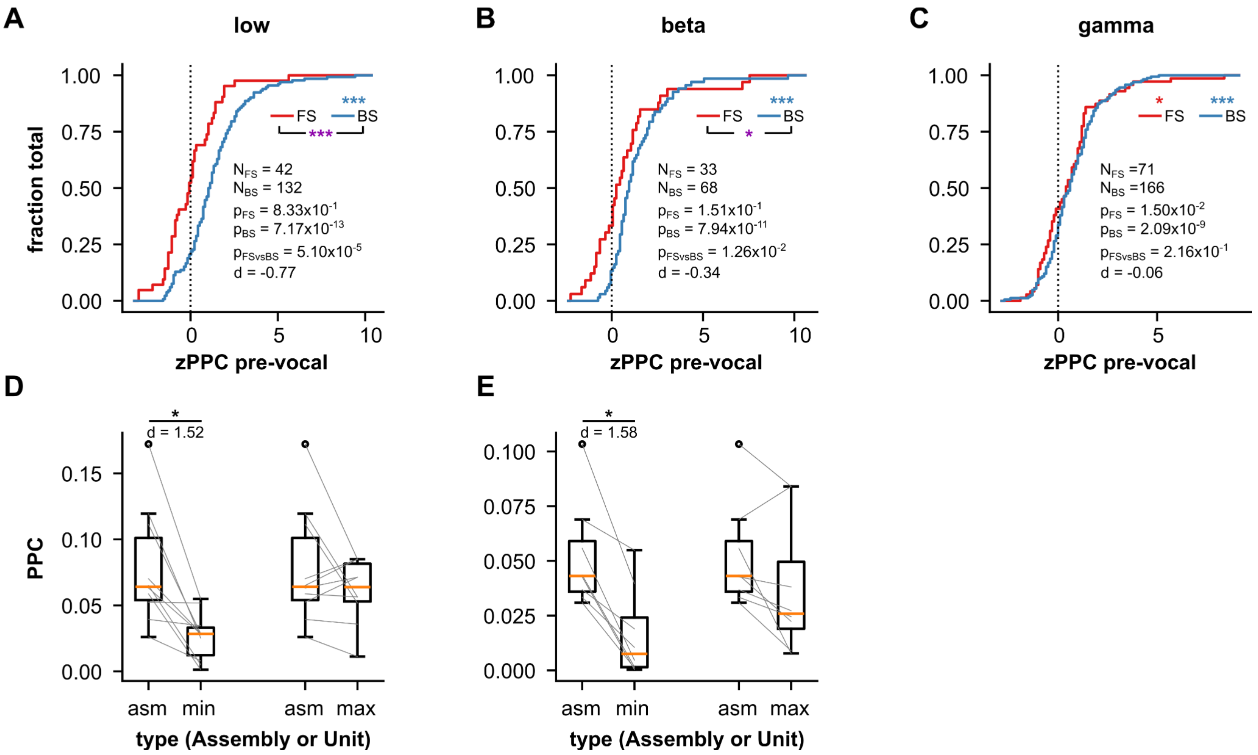


**Fig S9. Pairwise-phase consistency (PPC) in AC during pre-vocal periods.**

(**A**) Cumulative distributions of z-normalised (relative to no-voc periods) PPC values computed between pre-vocal spiking and low-frequency (2-10 Hz) LFPs, shown for significantly (Rayleigh test, $p < 0.05$) synchronized FS ($N_{FS} = 42$; red) and BS ($N_{BS} = 132$, blue) units during the pre-vocal period. Z-normalised PPC was not significantly different than 0 across FS units (FDR-corrected Wilcoxon signed-rank test, $p_{FS} = 0.83$), but was significantly above 0 for BS units ($p_{BS} = 7.17\text{x}10^{-13}$, blue ***). Normalised PPC values from FS units were significantly smaller than those from BS units (FDR-corrected Wilcoxon rank-sum test, $p_{FSvsBS} = 5.10\text{x}10^{-5}$, purple ***) with moderate effect size (d = -0.77). (**B**) Same as **A**, but spike-LFP synchrony was calculated using beta-band (12-20 Hz) LFPs. ($N_{FS} = 33$, $N_{BS} = 68$; $p_{FS} = 1.51\text{x}10^{-1}$, $p_{BS} = 7.94\text{x}10^{-11}$; $p_{FSvsBS} = 1.26\text{x}10^{-2}$, d = -0.34). (**C**) Same as **A**, but gamma-band LFPs were used for PPC calculation ($N_{FS} = 71$, $N_{BS} = 166$; $p_{FS} = 1.50\text{x}10^{-2}$, $p_{BS} = 2.09\text{x}10^{-9}$; $p_{FSvsBS} = 2.16\text{x}10^{-1}$, d = -0.06). Red star indicates FS z-normalised FS values significantly different from 0. (**D**) Comparison between assembly PPC values (asm) vs. PPC values of constituent units with the smallest PPC (*min*) or the highest PPC (*max*), with spike-LFP synchrony computed for low-frequency LFPs during pre-vocal periods. Data is shown only for assemblies significantly synchronised to low-frequency LFPs (Rayleigh test, $p < 0.05$; N = 10). Assembly PPC values were significantly higher than those for its constituent *min* unit (FDR-corrected Wilcoxon signed-rank test, $p_{corr} = 0.01$) with large effect size (d = 1.52), but not for those of its constituent *max* unit ($p_{corr} = 0.74$). (**E**) Same as in D, but values related to beta-band LFPs are shown (N = 8). Assembly PPC values were significantly higher than those for its constituent *min* unit ($p_{corr} = 0.02$) with large effect size (d = 1.58), but not for those of its constituent *max* unit ($p_{corr} = 0.54$). In the figure: *, $p_{corr} < 0.05$; ***, $p_{corr} < 0.001$. Assembly PPC in the gamma-range were not shown because the number of significant assemblies was low (N = 1).
